# Nonviral base editing of *KCNJ13* mutation preserves vision in an inherited retinal channelopathy

**DOI:** 10.1101/2022.07.12.499808

**Authors:** Meha Kabra, Pawan K. Shahi, Yuyuan Wang, Divya Sinha, Allison Spillane, Gregory A. Newby, Shivani Saxena, Yao Tong, Yu Chang, Amr A. Abdeen, Kimberly L. Edwards, Cole O. Theisen, David R. Liu, David M. Gamm, Shaoqin Gong, Krishanu Saha, Bikash R. Pattnaik

## Abstract

Clinical genome editing is emerging for rare disease treatment, but one of the major limitations is the targeted delivery of CRISPR editors. We delivered base editors to the retinal pigmented epithelium (RPE) in the mouse eye using silica nanocapsules (SNC) as a treatment for retinal degeneration. Leber Congenital Amaurosis (LCA16) is a rare pediatric blindness caused by point mutations in the *KCNJ13* gene, a loss-of-function inwardly rectifying potassium channel (Kir7.1) in the RPE. SNC carrying adenine base editor (ABE8e) mRNA and single-guide RNA precisely and efficiently corrected *KCNJ13^W53X/W53X^* mutation. Editing in both patient fibroblasts (47%) and human-induced pluripotent stem cell-derived RPE (LCA16-iPSC-RPE) (17%) had a negligible off-target response. Functional Kir7.1 channels were recorded from the edited LCA16-iPSC-RPE. In the LCA16 mouse model (*Kcnj13^W53X^*^/+ΔR^), RPE cells targeted SNC delivery of ABE8e mRNA preserved normal visual function measured by full-field electroretinogram (ERG). Moreover, multifocal ERG confirmed the topographic measure of electrical activity primarily originating from the edited retinal area at the injection site. Preserved retina structure, post-treatment, was established by Optical Coherence Tomography (OCT). This preclinical validation of targeted ion channel functional rescue, a challenge for pharmacological and genomic interventions, reinforces the effectiveness of nonviral genome editing therapy for rare inherited disorders.

**Graphical abstract:** 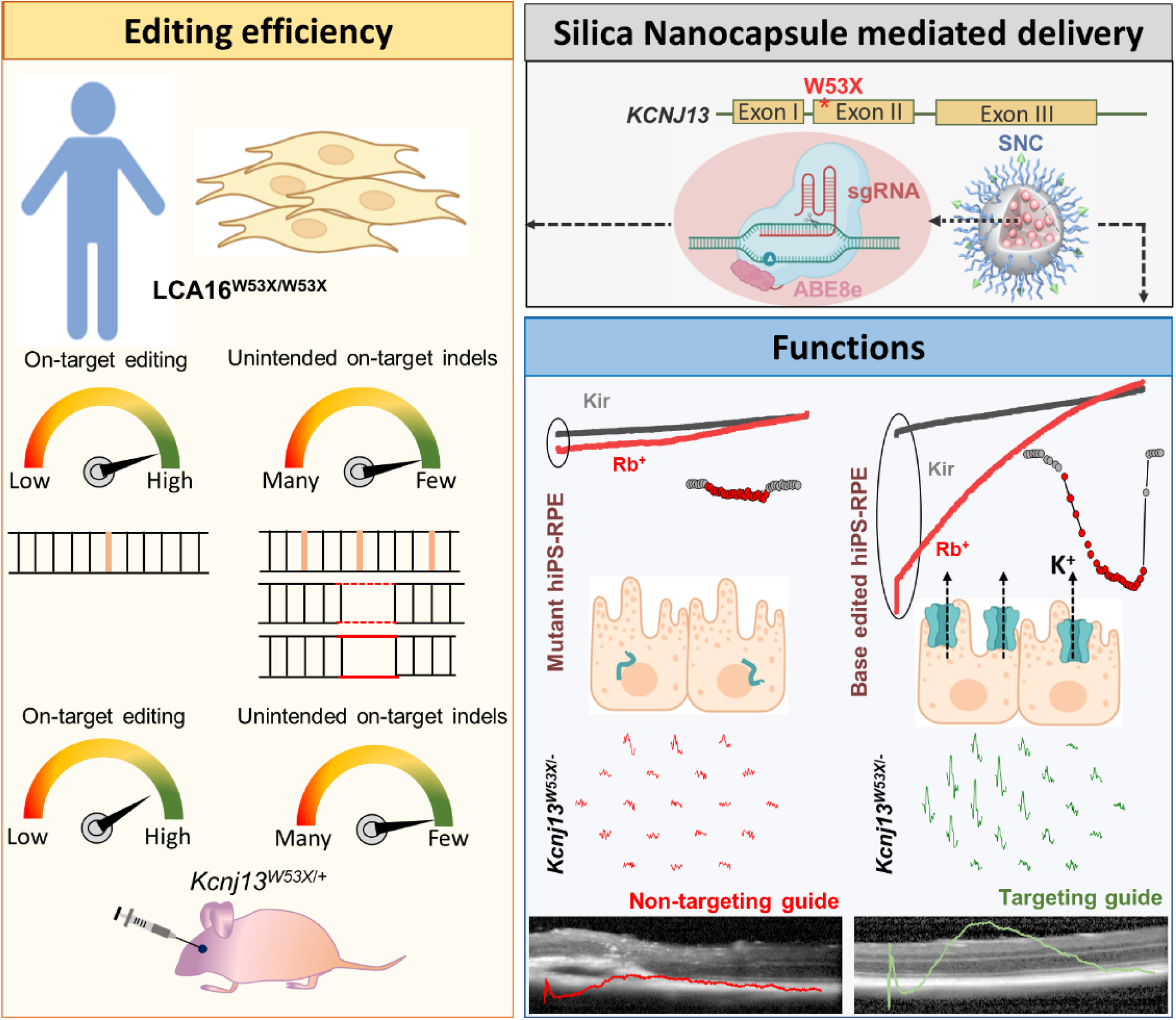

## Introduction

Leber congenital amaurosis (LCA16; OMIM#614186) is a severe autosomal recessive inherited retinal dystrophy (IRD) leading to blindness in early life. LCA16 is caused by the loss of function of the Kir7.1 potassium ion channel (encoded by the *KCNJ13* gene, location#2q37.1, OMIM#603208) in the apical processes of the retinal pigmented epithelium (RPE) cells of the eye (1, 2). The Kir7.1 channel controls K^+^ homeostasis in the subretinal space, supporting photoreceptor-RPE cell-cell signaling in support of visual functions such as phototransduction and phagocytosis. Several missense and nonsense loss-of-function mutations have been reported in *KCNJ13*, leading to diminished K+ conductance and altered electroretinogram (ERG) of the RPE (3–10).

There is no therapy available for LCA16 patients in the clinic. An LCA16 mouse model with biallelic mutations in *Kcnj13* does not survive long enough to test and develop possible therapeutic strategies. We previously used induced pluripotent stem cells (iPSCs) from a patient with the homozygous W53X mutation in *KCNJ13* to derive RPE cells (LCA16 iPSC-RPE) as a model to test potential treatments like translational readthrough-inducing drugs (TRIDs) and AAV-mediated gene augmentation (11). However, these possible therapeutics have limitations. TRID-mediated readthrough will broadly impact the proteome and may result in the introduction of non-functional amino acids at the site of readthrough. AAV-mediated gene augmentation does not correct the underlying mutations. It may yield only a transient effect alongside the innate and adaptive immune responses that diminish their therapeutic effect, particularly if repeated doses are required.

In contrast, gene editing could, in principle, correct the endogenous gene and permanently reverse the underlying cause of the disease. However, editing using Cas nucleases is limited by the inefficiency of precise editing via homology-directed repair (HDR) in most retinal cell types (12–14). Since HDR is inefficient in non-dividing cells such as the RPE, most edited cells will acquire new indel mutations, and cells with one corrected allele frequently co-inherit a disruptive indel allele. Moreover, double-strand DNA break (DSB) formation at the target site would pose a significant risk of uncontrolled indels and other undesired cellular consequences such as chromosomal translocations, large deletions, aneuploidy, p53 activation, chromothripsis, or transposon insertions (15–24). Base editing has the potential to overcome these limitations by capitalizing on the RNA-guided programmability of CRISPR-Cas9 to deliver a base modification enzyme that can site-specifically convert one single nucleotide to another (25).

Most base editors use a Cas9 nickase fused to an engineered or evolved deaminase enzyme (26–28). Adenine base editors (ABEs) install A>G changes, while cytosine base editors (CBEs) install C>T changes, both within a small editing window specified by the single guide RNA (sgRNAs) (29). Base editing does not rely on HDR and therefore tends to be efficient in non-dividing cells. While delivery of the larger base editor effectors can be challenging, nonviral silica nanocapsules (SNCs) provide significant advantages in that they can transiently and cell-specifically deliver a variety of payloads, including nucleic acids and proteins without integrating into the genome, with decent payload loading content (∼10 wt%) and encapsulation efficiency (>90 %) for all of the biologics mentioned above (30–32). We recently reported that SNCs decorated with all-*trans*-retinoic acid (ATRA) could achieve tissue-specific delivery to RPE cells via subretinal injection. This provides an unrealized opportunity to explore SNC-mediated base editor delivery for precise gene correction in LCA16 (33).

Here, we explored the ability of an SNC-delivered base editor to correct the endogenous pathogenic W53X *KCNJ13* point mutation in (1) patient-derived fibroblasts, (2) iPSC-derived RPE, (3) heterozygous knock-in *Kcnj13^W53X^*^/+^ mouse, and (4) wildtype allele disrupted *Kcnj13^W53X^*^/+ΔR^ mice via subretinal injection. ABE significantly outperformed a CRISPR-Cas9 HDR strategy in terms of efficiency and editing precision. SNC-mediated delivery was sufficient to achieve ∼16% editing of RPE cells in the mouse retina. Moreover, the high editing efficiency of 47% in patient-derived fibroblasts suggests the opportunity to pursue cell therapies using the engraftment of autologous edited and reprogrammed cells. We then demonstrate functional restoration of the Kir7.1 channel in patient-derived iPSC-RPE and in vivo in the LCA 16 mouse (*Kcnj13^W53X^*^/+ΔR^) model, further revealing the potential of CRISPR base editing to preserve vision in LCA16 patients. Our work provides essential proof-of-concept that base editing via SNC delivery is a viable strategy to prevent the progression of inherited retinal diseases, particularly channelopathies like LCA16.

## Results

### ABE8e mRNA efficiently corrects the HEK^W53X/W53X^cell line

Given the relatively low efficiency and heterogeneous outcomes of the traditional Cas9 nuclease-mediated gene editing approach (Supplementary Note and Supplementary Figure 1A-D & 2A-H), we focused on base editing to correct the W53X mutation. Base editors are engineered proteins that use the programmable DNA targeting ability of Cas9 to bring a nucleotide base-modifying enzyme to a precise editing window at the target DNA site. We used three bioinformatic tools for sgRNA design (see Methods). This leads to the targeted conversion of one or more bases, resulting in an altered DNA sequence without requiring DSBs or relying on cellular DNA repair machinery. Since the KCNJ13 causes the W53X mutation, c.158G>A nucleotide change, an ABE-mediated base editing strategy can potentially correct the mutation through an A>G conversion. To screen ABEs and validate the sgRNAs for their ability to edit this pathogenic locus with functional restoration of the Kir7.1 channel, we generated human embryonic kidney cells that stably express GFP-tagged KCNJ13-W53X (HEKW53X/W53X), and an isogenic WT control line, (HEKWT/WT) in which the WT KCNJ13 sequence was inserted into a single Flp-In recognition target (FRT) site at a transcriptionally active genomic locus (Figure 1A-B). We chose the ABE8e base editing effector due to its high efficiency (34, 35) and designed sgRNAs based on the predicted editing window at protospacer positions 4-8, counting PAM as position number 21-23. Only one of the three sgRNAs (G*C*G*CUAGCGUUGGAUGAUGU, PAM; TGG) was predicted to have high specificity while also placing the c158G>A position within the editing window of ABE8e (Figure 1C).

**Figure 1:**
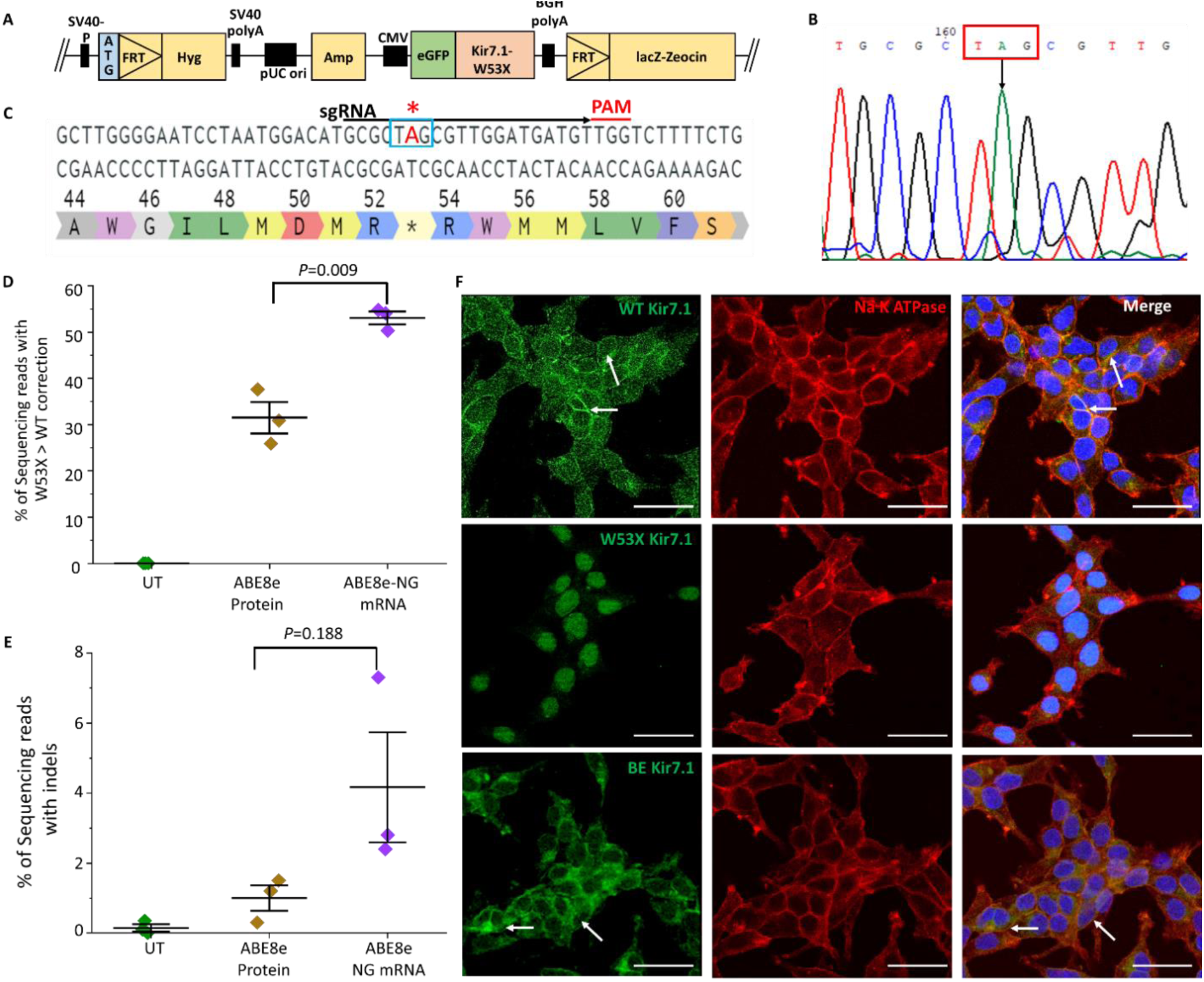
Evaluation of ABE8e RNP and ABE8e mRNA to correct hKCNJ13^W53X/W53X^ allele in HEK293 FRT stable cells. **(A)** Construct design to generate HEK293 FRT stable cells harboring the KCNJ13 ^W53X^ allele. **(B)** Schematic of the hKCNJ13 locus highlighting the mutation c.158G>A (blue box marked with *) and position of the W53X targeting sgRNA (black line) with TGG PAM (red line). **(C)** Chromatogram generated from HEK293 FRT stable cells showing the W53X codon marked in the red box and the downward black arrow indicating the specific nucleotide change (G>A). **(D)** Base editing efficiencies are shown as the percentage of sequencing reads with the corrected WT allele (and no other silent changes, bystander edits, or indels) in HEK293^W53X^ cells following electroporation of ABE8e protein + sgRNA (RNP) or ABE8e mRNA + sgRNA (n=3). Markers (diamonds) represent the individual biological replicates (n=3), and error bars represent SEM. **(E)** % of sequencing reads with indels in ABE8e RNP and ABE8e mRNA treated stable cells (n=3). Markers (diamonds) represent the individual biological replicates (n=3), and error bars represent SEM. **(F)** Kir7.1 expression in ABE8e mRNA treated cells assessed by immunocytochemistry. GFP primary antibody was used to enhance the endogenous signal. DAPI was used to stain the nucleus—scale bar, 50 μm. The white arrows mark the membrane localization in cells.

We observed ∽ 100% (99.7 ± 0.03%) transfection efficiency in HEK293 cells by electroporation of 3 µg of GFP mRNA (Supplementary Figure 3A-C). Therefore, we tested the activity of ABE8e in mRNA and protein formulations via electroporation. HEK^W53X^ cells were electroporated with either a mixture of ABE8e mRNA and sgRNA or ABE8e:sgRNA RNP complexes. Deep sequencing analysis was performed on the treated cells by isolating RNA instead of DNA to distinguish the inserted allele from the endogenous allele of the HEK genome. The pool of electroporated HEK^W53X^ cells showed significantly higher AT to GC correction efficiency with ABE8e mRNA (53.02 ± 1.38%) compared to ABE8e RNP (31.44 ± 3.39%) (Figure 1D). Our on-target analysis showed comparatively lower indels introduced using RNP (1.00 ± 0.36%) than mRNA (4.17± 1.57%) at and around the protospacer (Figure 1E).

The W53X nonsense mutation in *KCNJ13* disrupts its protein expression in HEK^W53X^ cells, so we assessed whether base editing restored Kir7.1 expression and subcellular localization at the protein level. Immunocytochemistry demonstrated that the Kir7.1 protein is present in the membrane in most of the base-edited HEK^W53X^ cells, similar to control HEK^WT^ cells. On the other hand, the fluorescence signal in untreated HEK^W53X^ cells is accumulated in the cytoplasm and nucleus due to its premature truncation (Figure 1F). These results confirmed the successful translation and trafficking of full-length protein after editing (Figure 1F).

To determine Kir7.1 channel function, whole-cell currents were recorded using an automated patch-clamp system from pools of either base-edited HEKW53X cells, untreated HEKW53X mutant cells, or untreated HEKWT cells (Figure 2A-C). In a standard external physiological HR solution, HEK^WT^ cells had a considerable negative membrane potential (−74.24 ± 4.16 mV, n=3 individual cells) (Supplementary Figure 4A). They exhibited an inward rectifying average Kir7.1 current of −45.0 ± 11.31 pA (n=62 individual cells) at −150 mV in 5 mM K^+^. The inward current increased in response to external 140 mM Rb^+^ (a known activator of the Kir7.1 channel) by 4.5-fold (−204.92 ± 37.33 pA) (Supplementary Figure 4B). However, the current was not blocked by adding 20 mM Cs^+^ (a known blocker of the Kir7.1 channel) to the bath solution (−41.50 ± 11.17 pA), suggesting a low Cs^+^ sensitivity of the Kir7.1 channel (Figure 2A) as reported earlier(1, 34). These responses were not observed in untreated HEK^W53X^ mutant cells, with a significantly lower resting current amplitude (−9.71 ± 1.26 pA) and negligible change upon adding Rb^+^ or Cs^+^ (Figure 2B). Importantly, base-edited HEK^W53X/W53X^ cells exhibited restored Kir7.1 function in 80% of cells (n=60), and the characteristic Rb^+^-responsive K^+^ current (−33.33 ± 7.68 pA baseline; −415.22 ± 54.0 pA with Rb^+^, −42.19 ± 9.56 pA with Cs^+^) (Figure 2C). The K^+^ current profile of 20% of the treated HEK^W53X^ cells (n=15) was comparable to mutant HEK^W53X/W53X^ untreated cells, which we attribute as unedited cells (K^+^ current in HR; −30.92 ± 8.04 pA, Rb^+^; −57.67 ± 12.46 pA, Cs^+^; −13.89 ± 5.07 pA) (Figure 2C). From these experiments, we conclude that ABE8e mRNA + sgRNA delivery can correct the W53X mutation at high efficiency (around 50% of total alleles and in 80% of cells), restoring Kir7.1 protein levels and K+ conductance.

**Figure 2:**
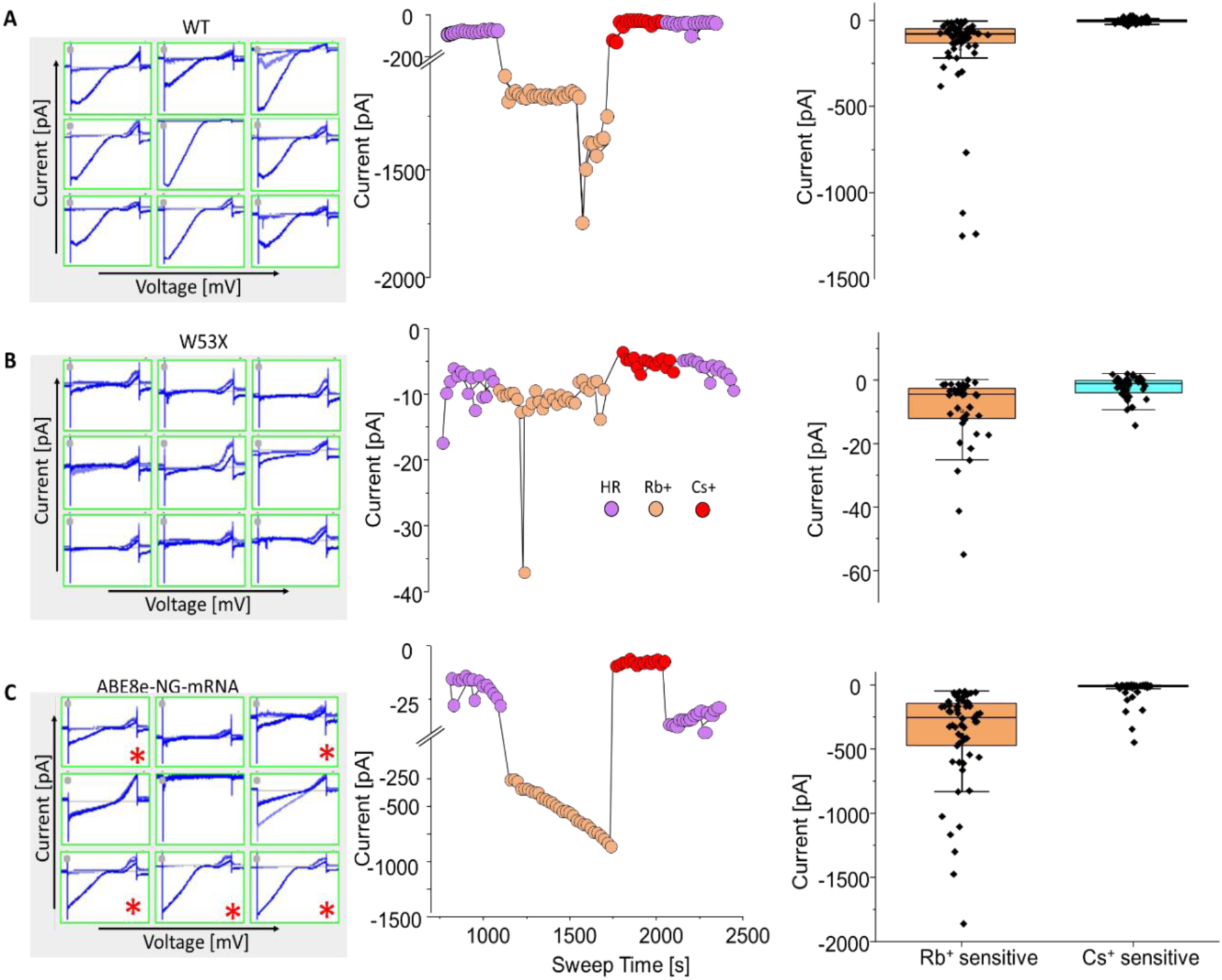
Kir7.1 current recording in WT, W53X, and base edited W53X HEK293 FRT stable cells. **(A)** Left snapshots of Kir7.1 current profile in WT stable cells. Center, the Current-Sweep plot represents the experimental timeline and is shown for one representative cell. Right, Rb^+^ and Cs^+^ sensitive current in HEK^WT^ stable cells **(B)** Left, snapshots of Kir7.1 current profile in HEK^W53X^ stable cells. Center, the Current-Sweep plot is shown for one representative cell. Right, Rb^+^ and Cs+ sensitive current in HEK^W53X^ stable cells. **(C)** Left, Snapshots of Kir7.1 current profile in HEK^W53X-BE^ cells using ABE8e mRNA. The cells marked with * showed recovery of K^+^ channel functions after the base editing. Center, the Current-Sweep plot is shown for one representative cell. Right, Rb^+^ and Cs+ sensitive current in HEK^W53-^cells.

### Efficient ex vivo base editing in patient-specific fibroblasts using nanoparticles

To explore the specificity of ABE at the endogenous human *KCNJ13* locus within patient-specific cells, we encapsulated ABE8e mRNA and sgRNA in SNC-PEG nanocapsules (Figure 3A) (33) for delivery into fibroblasts derived from an LCA16 (W53X homozygous) patient (Fibro^W53X/W53X^). Five days post-treatment, we assessed base editing efficiency using deep sequencing. As expected, the activity of ABE8e yielded efficient AT to GC editing (52.31 ± 0.06%) at the target W53X site (A^6^). Cells from both the treated and untreated populations (Supplementary Figure 5A-C) had low levels of other substitutions (or possibly sequencing error) in the amplicon, including bystander A>G and other T>G, T>C, G>A substitutions, with ABE8e exhibiting 6.48 ± 1.44% unwanted substitutions, and untreated Fibro^W53X/W53X^ cells showing 2.56 ± 0.02% unwanted substitutions (Figure 3B). Rare indel outcomes were detected in the ABE8e-treated samples (2.71 ± 0.40%) (Figure 3B). Bystander A>G editing near the target nucleotide was rare in the ABE8e-treated samples (A^14^; 0.26 ± 0.03%, A^17^; 0.07 ± 0.07%) (Figure 3C). We also marked some ABE activity upstream of the protospacer at positions A^-2^ (0.34 ± 0.025%) and A^-9^ (0.08 ± 0.08%) (Figure 3C). This bystander editing within and outside the protospacer resulted in silent (L48L) and missense (D50G, M51V, M56V, M57V) mutations (Figure 3D). Although these mutations were observed at a very low frequency (<1%) in Fibro^W53X/W53X^ (Figure 3E), it is possible to employ clonal selection of treated fibroblasts before iPSC generation and select a line that carries no bystander/incorrect edits.

**Figure 3:**
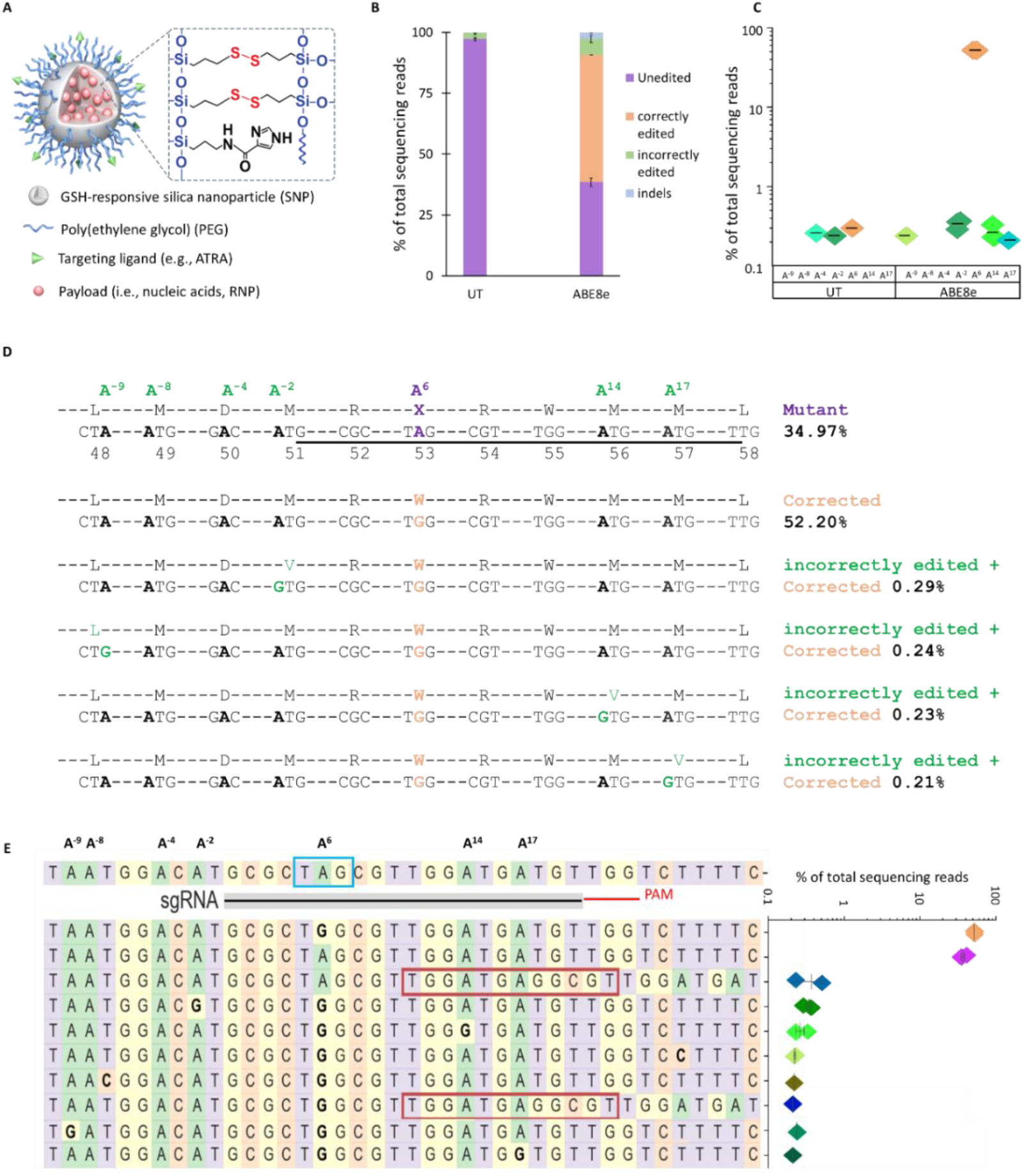
Evaluation of ABE mRNA + sgRNA combinations to correct the W53X allele in LCA16 patient fibroblasts. **(A)** Design of silica nanocapsules used to encapsulate ABE8e mRNA and sgRNA. **(B)** Base editing efficiencies are shown as the % of total DNA sequencing reads, classified as unedited, correctly edited, incorrectly edited due to bystander ‘A’ edits, and with indels in treated and untreated (UT) cells. **(C)** % Editing of the target (A^6^) and bystander (A^-9^, - A^-8^, A^-4^, A^-2^, A^14^, A^17^) ‘A’ to ‘G’ by ABE8e mRNA as observed in three independent experiments. **(D)** Amino acid conversion at the respective location was generated due to target and bystander edits. The protospacer sequence is underlined, the pathogenic early stop codon is in a purple box, the target ‘A>G’ edit is marked in orange and bystander ‘A’ edits are in green. **(E)** The sgRNA location is marked by a black line, PAM by a red line, and mutation in the blue box. All the ‘A’ bases within the protospacer are numbered 1-20 based on location. The ‘A’ bases downstream of the protospacer are numbered from −1 to −9, considering +1 as the first base of the protospacer. The top 10 most frequent alleles generated by ABE8e mRNA treatment show the nucleotide distribution around the cleavage site for sgRNA. Substitutions are highlighted in bold, insertions are shown in the red box, and dashes show deletions. The scatter plot shows the frequency of reads observed in treated cells (n=3 biological replicates). Figures presenting data from replicates are mean ± SEM.

In Fibro^W53X/W53X^, we also assessed the activity of another ABE mRNA (ABE7.10max), which has reduced activity and may reduce bystander mutations (Supplementary Figure 6A-C). This particular ABE mRNA could be advantageous in cases where fewer on-target corrections could have a beneficial outcome while avoiding detrimental bystander mutations. Unfortunately, the treated cells showed substantially lower AT to GC editing efficiency (14.93 ± 1.73%; p-value = 0.000027) than ABE8e at the target W53X site (A^6^). Undesired editing outcomes were also reduced: bystander editing at A^14^; 0.09 ± 0.09%, A^-2^; 0.07 ± 0.07%, A^-4^; 0.15 ± 0.08%, other observed substitutions (4.67 ± 0.79%) and indels (1.35 ± 0.42%) in the amplicon (Supplementary Figure 6A-C). These results confirmed that ABEs, particularly ABE8e, can efficiently revert the endogenous W53X nonsense mutation to wildtype in patient-derived fibroblasts.

### Restoration of Kir7.1 channel function in patient-derived iPSC-RPEs

Next, as a model for in vivo gene editing therapies for LCA16, we applied our base editing strategy to patient iPSC-derived RPE harboring homozygous W53X mutation in *KCNJ13*. Previous reports suggest that delivery into RPE by lipofection is challenging. Also, transient expression of BE machinery is preferred over long-term expression via AAVs to restrict off-target editing that may accumulate during prolonged exposure to genome editors. So, we used SNC-PEG as a delivery vector for ABE8e mRNA. We observed 27.44 ± 1.67% transfection efficiency in iPSC RPE cells using SNC-PEG (Supplementary Figure 7D). We did not observe any toxicity in the ABE8e-treated iPSC-RPE^W53X/W53X^, and cell morphology was comparable to untreated iPSC-RPE^W53X/W53X^ (Figure 4A). Deep sequencing analysis in the base-edited iPSC-RPE^W53X^ revealed 17.82 ± 1.53% editing of the mutant nucleotide to the wildtype sequence following ABE8e treatment (Figure 4B). Less than 4% of resultant alleles carried undesired genomic outcomes: indels quantified in treated RPE had a frequency of 0.68 ± 0.11% compared to 0.01 ± 0.002% in untreated controls, and bystander substitutions quantified in treated RPE totaled 2.94 ± 0.42% compared to 1.78 ± 0.07% in untreated controls (Figure 4C, Supplementary Figure 7A-C).

**Figure 4:**
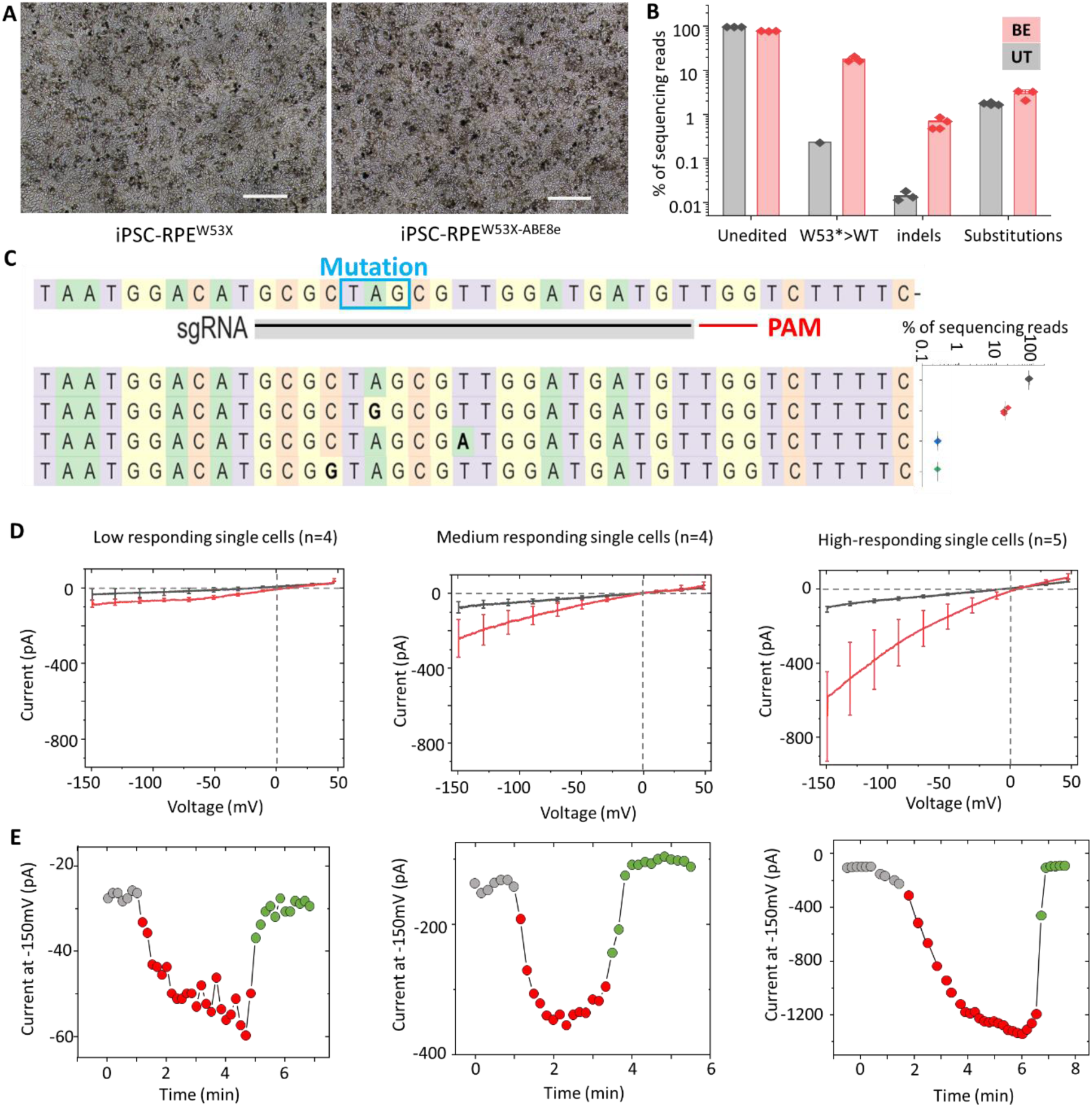
Evaluation of ABE8e to correct W53X alleles in iPSC-RPE^W53X/W53X^. **(A)** Representative bright-field images of base editor treated and untreated iPSC RPE^W53X/W53X^. Scale bars = 100 µm. **(B)** Base editing efficiencies following treatment (BE) with ABE8e mRNA and sgRNA encapsulated in SNC-PEG in iPSC-RPE^W53X/W53X^ as compared to untreated (UT) cells. Reads from the UT and treated cells (n=3) were categorized into 4 subtypes based on their sequences, unedited, W53*>WT, indels, and substitutions. **(C)** Reads generated by ABE8e mRNA treatment showing the nucleotide distribution around the cleavage site for sgRNA. Substitutions are highlighted in bold. The scatter plot shows the frequency of alleles observed in treated cells (n=3). Lines and error bars are represented as mean ± SEM. **(D)** Manual single-cell patch-clamp assays on iPSC-RPE^W53X^ cells after treatment with ABE8e. Of the 13 cells assessed for Kir7.1 activity, each could be binned into one of 3 classes: low-responding single cells, which appeared to be unedited mutant cells; medium-responding single cells, which showed a low level of Rb^+^ response; and high-responding single cells, which showed Rb^+^ response like iPSC-RPE^WT^ cells. The number (n) of cells binned into each class is shown at the top of each graph. **(E)** Current-sweep plot from a representative cell of each bin across a time course of being exposed to physiological HR solution (gray), Rb^+^ stimulation (red), and subsequent wash with HR solution (green).

Manual patch-clamp electrophysiology was carried out on the pool of base-edited iPSC RPE^W53X/W53X^ cells (Figure 4D) to assess the functional rescue of the Kir7.1 channel. Three types of Kir7.1 current profiles were observed in the edited iPSC-RPE^W53X/W53X^ pool. Some of the cells (n=5 out of 13) showed a normal amplitude of K^+^ conductance (−101.98 ± 0.07 pA) in HR solution, which was potentiated in Rb^+^ external solution by 8-fold (−820.97 ± 265.54 pA). These high-responding cells in the pool showed rescue of Kir7.1 channel function and were most likely to have only W53X>WT correction with or without neutral bystander mutations. Some of the cells did not show any recovery of Kir7.1 current (HR; −33.66 ± 31.59 pA, Rb^+^; −80.51 ± 19.37 pA) and had a similar profile to iPSC RPE^W53X/W53X^ cells (11). These low-responding cells (n=4 out of 13) likely experienced no base editing. A few cells (n = 4 out of 13) showed slightly higher Rb^+^-stimulated current than the untreated iPSC-RPE^W53X/W53X^ cells, but not as high as iPSC-RPE^WT/WT^ cells. The medium-responding cells showed only a 3-fold Rb^+^ response (−238.74 ± 102.13 pA) for the current amplitude observed in the HR solution (−72.57 ± 29.74 pA). We hypothesize that medium-responding cells contain a relatively low number of ion channels translated from a correctly base-edited *KCNJ13* allele. The current-sweep plots of the representative cell under different treatment solutions are shown in Figure 4E.

### *Kcnj13*^W53X/+^ monoallelic knock-in mouse as an LCA16 model for validating genome editing

A significant limitation to translating our *KCNJ13^W53X^* gene correction strategy to preclinical models is the lack of a mouse model harboring a homozygous pathogenic mutation in the gene. To test base editing efficiency in vivo, we first sought to create mice carrying a W53X substitution in the *Kcnj13* gene using CRISPR-Cas9 genome editing. In the pronuclei of zygotes, a combination of Cas9 protein, 2 different specific guides to the *Kcnj13* locus “(“GAATCCTAATGGACATGCGC<u>TGG</u>”” and ““TAATGGACATGCGCTGGCGC<u>TGG</u>””), and ssODN was injected (Figure 5A-B). Five of six newborn mice genotyped by sequencing had only one allele with the nucleotide alteration. This result was further validated by restriction fragment length polymorphism (RFLP) to digest the PCR product with the NheI enzyme, which selectively digests the mutant allele (Figure 5C). Breeding of heterozygous founders, and we determined that the homozygous mice, confirmed by genotyping (Figure 5D), resulted in postnatal day one lethality. The retinal structure (optical coherence tomography (OCT) images, Figure 5E) of newly generated heterozygous mice was identical to age-matched wildtype mice. Electrical activity generated by the retina was recorded using ERG as a measure of retinal function in response to a light stimulus. Specific retinal cells generate varying waveforms a-, b- and c-wave. The a-wave, a negative deflection, corresponds to the photoreceptor. The b-wave, which is positive, arises from the inner retinal cells, while the c-wave is generated by the RPE cell. Full-field electroretinogram (ERG) evaluation of heterozygous mice was identical to that of wildtype mice. (Supplementary Figure 8A-C), and c-wave amplitude measuring RPE cell function (W53X/+; 455.9 ± 18.7 vs. +/+; 504.5 ± 33.4 μV) were not statistically different (P=0.22) (Figure 5F).

**Figure 5:**
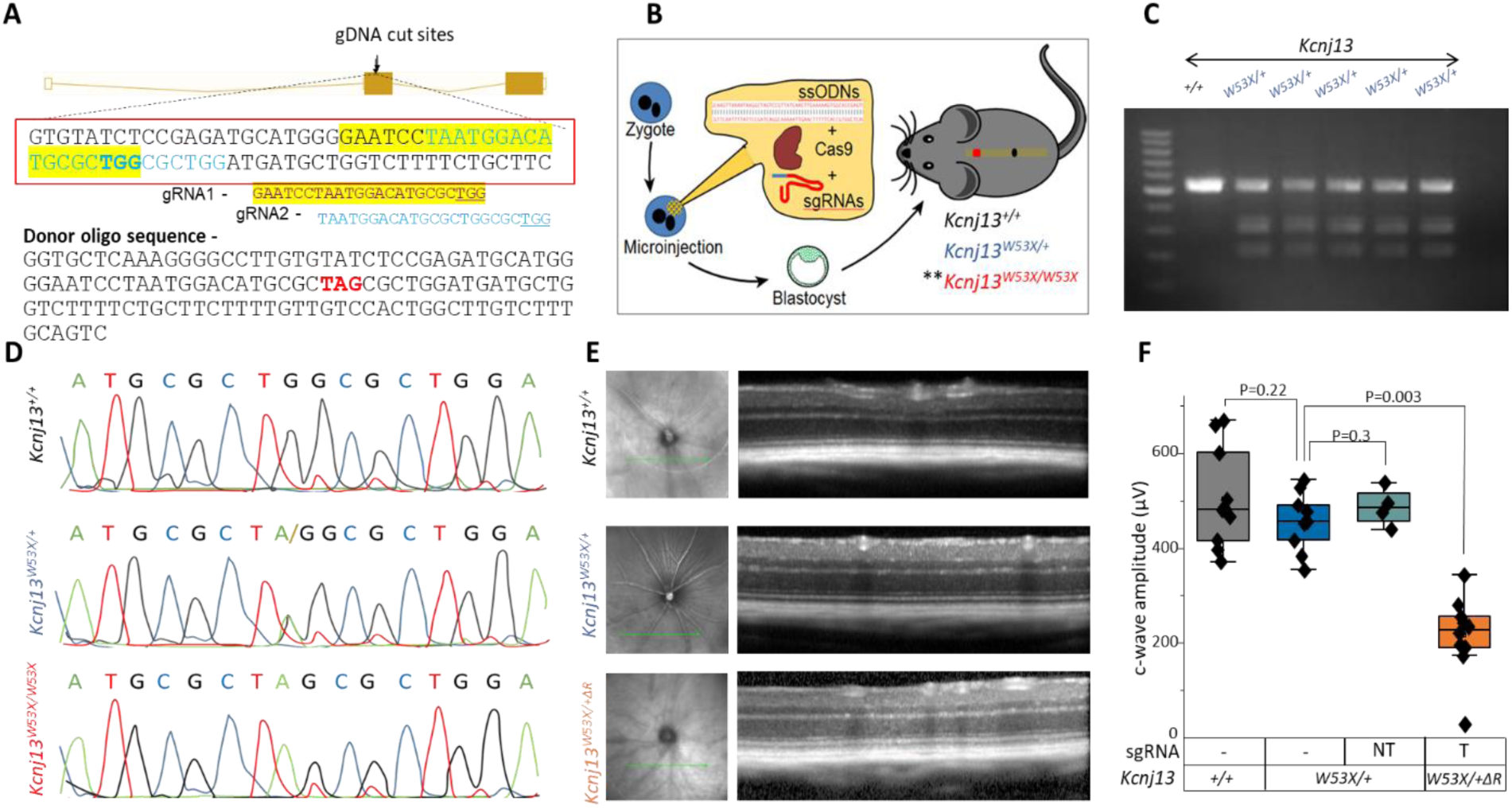
Visual function of *Kcnj13^W53X^*^/+^ mice is similar to *Kcnj13*^+/+^ mice. **(A** and **B)** Two different sgRNA targeting the *Kcnj13* gene at exon 2 and a ssODN sequence with the desired nucleotide change to generate the Kcnj13^W53X/W53X^ mouse model by CRISPR/Cas9 and HDR genome editing technique by microinjecting them into the pronuclei of the zygote. **(C)** RFLP analysis of the Kcnj13 gene from the newly generated mice digested with Nhe1 enzyme on 2% agarose gel. **(D)** Chromatograph confirming the mouse genotype. **(E)** OCT images showing the comparison between Kcnj13^+/+^, Kcnj13^W53X/+^, and WT allele-disrupted Kcnj13^W53X/+ΔR^ mice. **(F)** Averaged c-wave response confirming WT allele disruption in the RPE of Kcnj13^W53X/+ΔR^ using the targeted guide (T). (**= postnatal day1 lethal; NT= not targeting sgRNA)

Unlike humans, we have determined that at least one normal mouse *Kcnj13* gene allele is required for survival beyond postnatal day 1 (7, 35), precluding the establishment of a genetic model recapitulating the human recessive disease. Therefore, we used our *Kcnj13^W53X/+^* mice to generate an LCA16 mouse model (*Kcnj13^W53X/+ΔR^*) using a Cas9 nuclease complex targeted to disrupt the wildtype allele (Figure 5E-F). The wildtype allele was edited in the RPE cells by delivering Cas9 mRNA and guide through sub-retinal delivery of SNCs decorated with ATRA (SNC-PEG-ATRA), which explicitly targets RPE cells. The size and zeta-potential of ABE mRNA+sgRNA-encapsulated SNC were summarized in supplementary Table 1, and supplemental Figure 9. Mice were evaluated after 6 weeks to demonstrate intact retinal structure (Figure 6E, lower panel). ERG measurements of c-wave amplitude before the injection was 515.5 ± 9.5 μV, but 6 weeks after the disruption, it dropped to 234.2 ± 14.9 µV (P=0.003). A decrease in c-wave amplitude by 54 % reflects the functional loss of RPE cells as in LCA16. This disease model was used for the functional validation of base editing in vivo.

**Figure 6:**
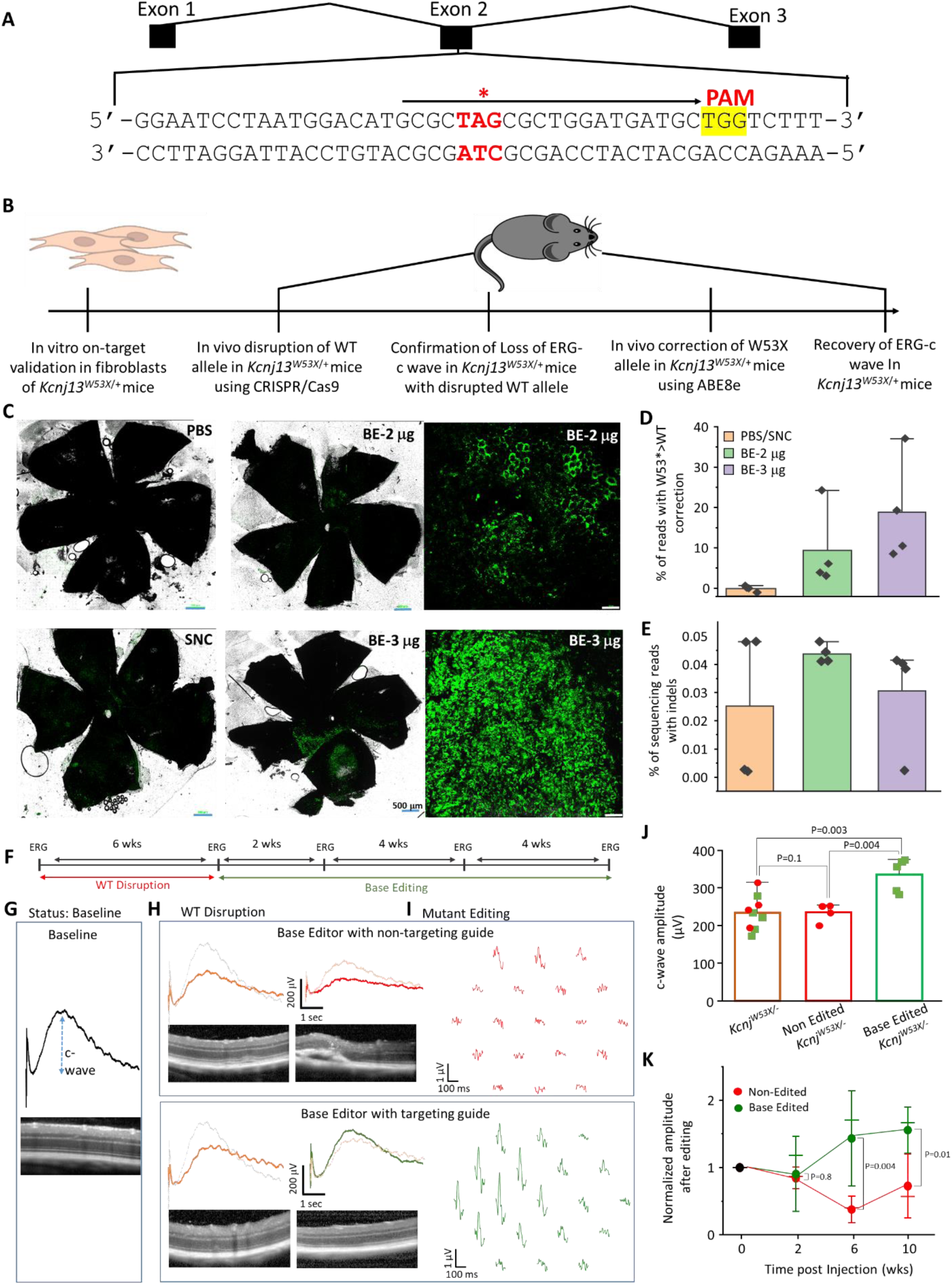
Phenotypic reversal of RPE^W53X^ mice following in vivo ABE8e treatment. **(A)** The sgRNA design targeting the Kcnj13^W53X^ allele for base editing, with the black arrow representing the sgRNA spacer sequence, the desired base editing site indicated by an asterisk, and the PAM shown in yellow. **(B)** Workflow of in vivo base editing strategy. **(C)** RPE floret of eyes subretinally injected with SNC-PEG-ATRA packaged ABE8e mRNA, W53X-sgRNA, and GFP mRNA or empty SNC-PEG-ATRA/PBS as a mock treatment. **(D)** % of cells with W53X>WT corrections, observed in *Kcnj13^W53X^*^/-^ mice treated with 2 μg or 3 μg of ABE8e. **(E)** % of sequencing reads showing the indels, observed in *Kcnj13^W53X^*^/-^ mice treated with 2 μg or 3 μg of ABE8e. **(F)** A timeline outlining the in vivo experiment. Baseline ERG is performed prior to the disruption of the WT allele and is followed up after 6 weeks. ERG is performed on mice prior to injection of the base editor, and recovery is monitored for 10 weeks.**(G)** Representation of the c-wave amplitude acquired from the *Kcnj13^W53X^*^/+^ mice with the OCT image of the retina. **(H)** the reduction in c-wave amplitude of the *Kcnj13^W53X^*^/+^ mice at 6 weeks after disrupting the wildtype allele with Cas9 protein and sgRNA guide specific to the wildtype allele. **(I)** c-wave traces and mfERG response following the injection of base editor with a non-targeting guide to the mutant allele (red) and base editor with a targeting guide (green). The faded traces are for comparison before the disruption of the WT allele (gray) and injection of the base editor (orange). **(J)** Bar graph comparing the c-wave amplitude 6 weeks after the disruption of the wildtype allele (blue) to the c-wave amplitude 6 weeks after the injection of base editor with a non-targeting guide (red) and targeting guide (green). **(K)** Normalized c-wave amplitude comparison between the eyes injected with non-targeting and targeting guides at weeks 2, 6, and 10. (ns= not significant).

### In vivo base editing via subretinal injection of ABE8e encapsulated SNCs

We designed sgRNAs targeting the W53X allele (Figure 6A) of mouse *Kcnj13*. We validated it in the fibroblasts derived from *Kcnj13^W53X/+^*mice via electroporation (Figure 6B). To determine the editing efficiency in a heterozygous model system (mouse fibroblasts and in vivo), we delivered using SNC-PEG. We calculated the increase in the WT reads by converting the available W53X allele. We observed a 14.5% increase in the WT reads in the treated fibroblasts suggesting 29% of cells received an edit (Supplementary Figure 10).

For in vivo subretinal delivery into *Kcnj13^W53X/+^* mice, ABE8e mRNA, sgRNA, and GFP mRNA were packaged into SNC-PEG-ATRA. The injection site could be located through GFP fluorescence (Figure 6C), GFP-positive RPE tissue, and the choroid optic cup were dissected for deep sequencing of the *Kcnj13* locus. We found that delivering 3 μg ABE8e mRNA resulted in 16.80 ± 7.87% RPE editing while delivering 2 μg resulted in 9.45 ± 5.02% RPE editing (n=4 eyes for each dose) (Figure 6D). Our on-target analysis showed no other A>G substitutions outside or within the protospacer region. In addition, indel mutations were not significantly increased over placebo-treated mice (Figure 6E). These data demonstrated the specific editing of the W53X allele in vivo by ABE8e-encapsulated SNC-PEG-ATRA.

### Base editing of the LCA16 mutant allele improves RPE function

In our *Kcnj13*^W53X/+ΔR^ mice, we edited the mutant allele using the base editor and specific guide packaged in SNC-PEG-ATRA and were administered subretinally. Full-field ERG measurements were performed at weeks 2, 6, and 10 to monitor longitudinal rescue in phenotype (Figure 6F). At week 10, mfERG was performed on injected eyes to determine if functional recovery was regionalized based on localized delivery of the base editor complex. For the control group, *Kcnj13*^W53X/+*ΔR*^ was injected with a base editor and a non-targeting guide.

The non-targeted guide-injected mice exhibited a progressive decrease in c-wave amplitude as well as depressed mfERG waveforms in almost the entire retina. On the other hand, the base-edited eyes showed an increase in the c-wave amplitude at all measured time points (Figure 6G-J). mfERG showed localized waveforms (Figure 6H and Supplementary Figure 11A) that perhaps correlate with the site of injection (Figure 6C). On the second week following the injection, the normalized c-wave amplitude (to that before the therapy) in eyes receiving the non-targeted guide was 0.84 ± 0.06, identical to the eyes injected with the targeted guide, 0.9 ± 0.18 (P=0.8). At the ten weeks post-injection, the c-wave was reduced (0.73 ± 0.19) for the non-targeted retina compared to an increasing trend (1.55 ± 0.11, P=0.01) for the targeted guide-delivered retina. This increase of 55% in amplitude and intact retinal structure, as evaluated by OCT (Figure 6H, 6J), demonstrates that inherited retinal channelopathies could be mitigated using base editors. As a further preclinical validation, we determined if the SNC used to deliver the base editor triggers an immune response. The RPE cells in culture were either transduced with lentivirus, AAV7m8, or AAV5 or were incubated with silica nanocapsule. Expression of inflammatory cytokines, including interferon-gamma (IFN-γ), interleukin (IL)-1α, IL-1β, IL-6, CD-8, and Iba1 was assessed by qPCR. CD8 and Iba1 showed increased expression in virally transduced RPE cells but not in cells exposed to SNC (Supplementary Figure 11B).

### Off-target analysis of sgRNA for human and murine W53X targets confirms the specificity

The Cas9 effector of the ABE dictates the specificity of the base editor to its target site and can bind off-target sites in the genome based on a similarity to the guide sequence. We used an online computational algorithm, Cas-OFFinder (36), to identify the putative off-target sites for the human W53X sgRNA used above in our ABE treatments. We found 2727 potential off-target sites, most harboring 4 mismatches with 1 nucleotide RNA bulge (n=2004) or DNA bulge (n=512), with other configurations and closer matches occurring much less frequently (Supplementary Figure 12). The likelihood of off-target editing should be greatest at sites with 1-2 mismatches and no DNA/RNA bulge, but our in-silico analysis showed no such off-target sites for this human W53X sgRNA.

We analyzed the off-target activity of ABE8e in base-edited HEK^W53X/W53X^, LCA^W53X/W53X^, and iPSC-RPE^W53X/W53X^ cells at the nine potential sites (supplementary Table 5). Deep sequencing analysis of these sites showed that ABE8e treatment did not lead to significant off-target A>G editing or introduction of indels compared to untreated control cells except at one locus. In particular, this one-off-target (Off_3: GRCh38:50530925 Intergenic) showed 4x more substitutions in base-edited HEK^W53X/W53X^ cells. However, this nucleotide is already reported in humans to have a natural A>G single nucleotide polymorphism (rs139941105, Minor Allele Frequency in 1000 Genomes=0.01) of no known clinical significance (Supplementary Figure 13A-B). These data demonstrate that the off-target modification levels of ABE8e at each of the nine potential sites are much lower than its on-target editing efficiency and overall indicate a favorable off-target editing profile.

To detect in vivo off-target editing, we used gDNA isolated from the optic cup of base editor-treated heterozygous *Kcnj13^W53X^*^/+^ mice and screened the 11 top predicted off-target sites by rhAMP-seq (supplementary Table 7). ABE8e treatment introduced on-target W53X>WT correction with an efficiency of 9.5% in these mice. Similar to our in vitro off-target analyses described above, we detected no substantial off-target editing following in vivo editing of mouse RPE (Supplementary Figure 13C-D). Together, these results suggest that ABE8e could be an effective and specific method to correct polymorphisms in the genome and that SNCs can serve as effective vehicles to correct mutations in the RPE.

## Discussion

Currently, recessive loss-of-function *KCNJ13* point mutations are the only known genetic cause of LCA16, disrupting the Kir7.1 channel function and altering RPE physiology, leading to retinal degeneration with progressive vision loss in patients. In the absence of disease-modifying approved therapies, correction of gene mutations in the RPE cells could avoid treatment-related serious adverse events (no off-target responses), restore the Kir7.1 channel function (in vitro and in vivo electrophysiology outcomes), improve vision, or prevent further disease progression (ERG and OCT). Of the 12 different known LCA16-causing *KCNJ13* gene mutations (37), 8 are theoretically accessible by base editors, and 5 of those by ABE (requiring A>G (or T>C) edits). Here, we provide essential proof-of-concept that delivery of ABE to RPE via subretinal injection of SNCs can result in correction of the W53X pathogenic allele and restoration of Kir7.1 channel function in RPE cells.

The SNC is a powerful nonviral delivery system for protein and nucleic acids (33). Our group has previously reported the SNC-PEG mediated transfection (50% for plasmid and 60% for mRNA), delivery, and GSH-responsive release characteristics (33, 38). The mRNA transfection efficiency remained unaffected at GSH concentrations lower than 0.5 × 10^−3^ M, suggesting extracellular stability of SNC-PEG (considering plasma/extracellular GSH concentration ranges from 10^−6^M to 2× 10^−5^ M). However, at GSH concentrations of 0.5 × 10^−3^ M or higher (corresponding to cytosol GSH concentrations between 10^−3^ M to 10^−2^ M), a significant decrease in the mRNA transfection efficiency was observed, indicating the SNC-PEG can effectively break down in the cytosol to release the payload readily. After treating the cells with the ABE8e mRNA loaded SNC-PEG, the nanoparticles will stay intact until internalized by the cells. Subsequent to uptake, intracellular GSH will facilitate nanoparticle disassembly and release the encapsulated ABE8e mRNA, enabling the targeted base editing within the iPSC-RPE cells. We observed >50% on-target editing in patient-derived fibroblasts, and 18% in iPSC-RPE cells using SNC-PEG delivery system. For in vivo editing of mouse RPE, we used the ligand-conjugated SNC-PEG-ATRA and observed 16% of RPE cells were edited. The editing of the RPE cells in vivo is confined, a result of targeted subretinal SNCs delivery to localized cells around the injection site. We previously showed that the pathological state of RPE can be reversed by rescuing 25% of the Kir7.1 channel function (11). A recent trial from Editas (Edit101) introduced indels in the *CEP290* gene linked to a different type of Leber Congenital Amaurosis, LCA10. It restored visual function in non-human primates with >10% functional photoreceptors (39). Each RPE sub serves multiple photoreceptors, so 10% RPE correction will rescue a significant multiple of these photoreceptors. This potential multiplicative effect makes the RPE even more valuable as a target for genome editing. And, like many other ocular diseases, treating LCA16 should not require the correction of all mutant cells. Therefore, the base editing efficiencies we gained in this study are well within the range of providing considerable therapeutic benefit.

Our study demonstrates an important new nonviral delivery method for gene correction in vivo with transient exposure to editing reagents. The ABE activity is anticipated to be transient within cells and on the order of a few hours, considering the half-life of Cas9 mRNA (15 mins −3 hours) and the half-life of the sgRNA (<3 hours) (40). Based on prior studies with transient Cas9 activity, we expect the off-target potential of our nanoparticle treatment to be lower than viral based delivery strategies (41). SNC-mediated base editing can restore Kir7.1 channel function, as assessed in our edited iPSC-RPE cells and in vivo *Kcnj13^W53X/+ΔR^* mice studies. Previous studies have demonstrated the functional outcomes of base editing in RPE by correcting a point mutation in the RPE65 enzyme. It restored its enzymatic substrates to support the visual cycle when rescued. Although ion channels such as Kir7.1 catalyze transmembrane flux of ions, their functional restoration has a less established record of success (42). On-target bystander substitutions and indels in some iPSC-RPE and in vivo mouse RPE cells are detectable. Still, they do not interfere with the potential functional benefits of repairing the mutation in the tissue as these genes are not expressed in the retina. Due to lacking a homozygous LCA16 mouse model, we created our novel in vivo model by disrupting the WT allele of RPE cells using subretinal delivery of SNC carrying the targeted guide. Our study establishes that in vivo editing with nonviral delivery systems can restore the Kir7.1 channel function in RPE. The edited *Kcnj13^W53X/+ΔR^* mice displayed recovery of their ERG c-wave amplitudes from the edited RPE and had no further retinal degeneration. Additional preclinical validation would require determining safety and efficacy using an acceptable animal model for regulatory clearance.

A limitation of our study is that we do not yet know the clonal editing outcomes of cells edited in vitro or in vivo. The combination of alleles in a single cell is a critical parameter for the function of the tetrameric Kir7.1 channel. Correction of just a single allele, while having detrimental edits in fellow alleles, will likely not preserve RPE health to a similar extent as a biallelic correction or monoallelic correction without alteration in the second allele. A clonal analysis would reveal the precise editing outcomes per cell. It could distinguish the phenotypic impact of edited genotypes (43). However, RPE cells from iPSC-RPE and mouse eyes are post-mitotic, making clonal amplification of edited cells experimentally intractable in these systems.

In conclusion, while addressing several significant challenges in correcting an LCA16-causing pathogenic mutation in the Kir7.1 channel, our study provides proof-of-concept therapy for a rare disease. Importantly, base editing of the *KCNJ13^W53X^* allele in vitro and in vivo showed specificity for W53X mutation without generating detrimental on-target bystander substitutions, indels, or off-target edits elsewhere in the genome. K^+^ conductance in iPSC-RPE in vitro and c-wave recovery in *Kcnj13^W53X/^*^-^ in vivo confirmed the functional rescue of the Kir7.1 channel following base editing. The specific delivery of base editor tools to RPE via the nonviral SNC platform provides a powerful emerging tool for tissue-specific delivery of mRNA or protein-based gene editing therapeutics with good biocompatibility, the ability to package large cargo, increased safety profile, and significantly streamlined manufacturing. These advances will likely direct future preclinical and clinical applications of base editing for correcting mutations causing LCA16 and other ocular genetic diseases.

## Methods

### HEK Flp-In™ 293 stable cells with GFP-tagged WT and W53X Kir7.1 expression

HEK Flp-In™ 293 host cells (ThermoFisher Scientific#R75007, MA) were generated using a pFRT/lacZeo target site vector to express GFP-tagged Kir7.1 (WT and W53X). These cells contain a single Flp Recombination Target (FRT) site at a transcriptionally active genomic locus to allow stable integration of GFP-tagged human *KCNJ13* sequence (WT and W53X). As these cells express the Zeocin^TM^ gene under SV40 early promoter, a complete D-MEM high glucose media (10% FBS, 1% Pen-Strep, 2mM L-glutamine) containing 100 µg/ml Zeocin^TM^ was used for maintenance. GFP-WT or GFP-W53X *hKCNJ13* gene sequence was integrated into the genome of these cells based on the manufacturer’s guidelines. Briefly, the cells were co-transfected with FLP-In™ expression vector (pcDNA5/FRT) containing GFP-tagged *hKCNJ13* sequence (WT or W53X) created by in-fusion cloning, and pOG44 recombinase expression plasmid. The pOG44 plasmid with constitutive expression of the Flp recombinase under CMV promoter mediates the homologous recombination between the FRT sites of host cells and the expression vector such that the construct from the vector is inserted into the genome at the FRT site. This insertion brings the Hygromycin B resistance gene into the frame, inactivates the zeocin fusion gene, and expresses the gene of interest under the CMV promoter. 48 hours post-co-transfection, the cells were passaged at 25% confluency to select stable transfectants in 200 µg/ml of Hygromycin B. The Hygromycin B resistant cell clones (n=15-20) were picked, maintained in 100 µg/ml Hygromycin B, and further expanded for their characterization by genotyping (Sanger sequencing) and protein expression (immunocytochemistry). The primers used for in-fusion cloning, and Sanger sequencing are listed in supplementary Table 2.

### Patient-derived fibroblasts and iPSC-RPE cell culture and maintenance

Fibroblasts derived from a skin biopsy of an LCA16 patient (7, 11) with homozygous W53X mutation in *KCNJ13* were cultured and maintained in complete DMEM high glucose media containing 10% FBS and 1% Pen-Strep at 37 °C with 5% CO_2_. Induced pluripotent stem cells (iPSCs), reprogrammed from patient-derived fibroblasts, were cultured on Matrigel and differentiated to iPSC-RPE using an approach similar to that previously described (Shahi et al. AJHG 2019; Sinha et al., AJHG 2020). Briefly, on D0 of differentiation, iPSCs were lifted using ReLeSR (Stem Cell Technologies; Cat# 05872) to generate embryoid bodies (EBs). The EBs were maintained overnight in mTeSR+ containing 10 µM ROCK inhibitor (R&D Systems; Cat# Y-27632). Then, over the next three days, the EBs were gradually transitioned to Neural Induction Media (NIM; DMEM: F12 1:1, 1% N2 supplement, 1x MEM nonessential amino acids (MEM NEAA), 1x GlutaMAX, and 2 μg/ml heparin (Sigma)). On D7, EBs were plated on Nunc 6-well plates coated with Laminin (ThermoFisher Scientific; Cat# 23017015; diluted 1:20 in DMEM/F12) on D16, neurospheres were mechanically lifted. The remaining adherent cells were transitioned to retinal differentiation media (RDM; DMEM/F12 (3:1), 2% B27 without retinoic acid, 1% antibiotic-antimycotic solution) and allowed to differentiate to RPE. RDM was supplemented with 10 µM SU5402 (Sigma-Aldrich; Cat# SML0443-5MG) for the first four media changes and 3 µM CHIR99021 (BioGems; Cat# 2520691). After >60 days of differentiation, iPSC-RPE cells were purified as described by Sharma et al. and cultured on the desired surface (44). Briefly, cultures with differentiated patches of iPSC-RPE were dissociated using 1X TrypLE Select Enzyme (ThermoFisher; Cat# 12563011) and enriched for iPSC-RPE via magnetically activated cell sorting using anti-CD24 and anti-CD56 antibodies (Miltenyi Biotec). iPSC-RPE cells in the negative cell fraction were seeded on the desired surface pre-coated with Laminin (ThermoFisher Scientific; Cat# 23017015; diluted 1:20 in DMEM/F12) and cultured.

### Silica nanocapsules (SNCs) for adenine base editing and CRISPR-Cas9 gene editing

We recently reported a safe and efficient nanoplatform, SNC, to deliver CRISPR gene editing and base editing components (33). SNCs were synthesized using a water-in-oil microemulsion method. The oil phase (1 ml) was prepared by mixing Triton X-100 (885 μL) with hexanol (0.9 mL) and cyclohexane (3.75 mL). An aliquot of aqueous solution (25 μL) containing the desired payload (ssODN or base editor mRNA + sgRNA, with a total nucleic acid concentration of 2 mg/mL) was mixed with the silica reagents: TEOS (4 μL), BTPD (6 μL) and N-(3-(triethoxysilyl) propyl)-1H-imidazole-4-carboxamide (TESPIC, 1 mg). The synthesis of TESPIC was reported previously (33). This mixture was homogenized by pipetting and then added to the oil phase (1 mL). The water-in-oil microemulsion was formed by vortexing for 1 min. Under vigorous stirring (1,500 rpm), an aliquot of 30% aqueous ammonia solution (4 μL) was added, and the water-in-oil microemulsion was stirred at 4 °C for 12 h to obtain unmodified SNCs. Acetone (1.5 mL) was added to the microemulsion to precipitate the SNCs. The precipitate was recovered by centrifugation and washed twice with ethanol and three times with ultrapure water. The purified SNCs were collected by centrifugation. The as-prepared, unmodified SNC was re-dispersed in ultrapure water (1 mL). For surface modification, mPEG-silane or a mixture of mPEG-silane + silane-PEG-NH_2_ (molar ratio of mPEG-silane: silane-PEG-NH_2_ = 8:2) was added to the SNC mentioned above suspension, for the synthesis of SNC-PEG without ATRA (i.e., SNC-PEG) or SNC-PEG-NH_2_ (for ATRA conjugation), respectively. The total amount of PEG is 10 wt% of SNC. The pH of the suspension was adjusted to 8.0 using a 30% aqueous ammonia solution. The mixture was stirred at room temperature for 4 h. The resulting SNCs were purified by washing with ultrapure water three times and concentrated with Amicon® Ultra Centrifugal Filters (Millipore Sigma, USA). ATRA was conjugated onto SNC-PEG-NH_2_ via EDC/NHS catalyzed amidation. Payload-encapsulated SNC-PEG-NH_2_ (0.5 mg) was re-dispersed in 1 mL DI water. EDC (7.5 μg), NHS (4.5 μg), and a DMSO solution of ATRA (6 μg in 5 μL DMSO) were added to the above solution. The solution was stirred at room temperature for 6 h, and then the resulting ATRA-conjugated SNC (i.e., SNC-PEG-ATRA) was washed with water three times. The SNC-PEG-ATRA was concentrated with Amicon Ultra Centrifugal Filters to a payload concentration of 2 mg/ml before use. **Materials:** Tetraethyl orthosilicate (TEOS), Triton X-100, acetone, ethanol, 1-ethyl-3-(3-dimethylaminopropyl)carbodiimide (EDC), N-hydroxysuccinimide (NHS), and ammonia (30% water) were purchased from Fisher Scientific, USA. Hexanol and cyclohexane were purchased from Tokyo Chemical Industry Co., Ltd., USA. Dimethyl sulfoxide (DMSO) was purchased from Alfa Aesar, USA. Bis(3-(triethoxysilyl)propyl)-disulfide (BTPD) was purchased from Gelest, Inc., USA. Methoxy-poly (ethylene glycol)-silane (mPEG-silane, Mn = 5000) and amine-poly (ethylene glycol)-silane (NH2-PEG-silane, Mn = 5000) were purchased from Biochempeg Scientific Inc., USA. All-trans-retinoic acid (ATRA) was purchased from Santa Cruz Biotechnology, USA.

### Base editor mRNAs

Using their mammalian-optimized UTR sequences, base editor mRNA was obtained as a custom product from Trilink Biotechnologies. The mRNAs were synthesized with complete substitution of uracil by N1-methylpseudouridine, co-transcriptional 5’ capping with the CleanCap AG analog resulting in a 5’ Cap1 structure, and included a 120 nucleotide polyA tail.

### sgRNA design

The sgRNAs targeting the W53X location in the human *KCNJ13* gene were designed using Benchling (https://www.benchling.com). The design was validated with two other online tools, CRISPR-RGEN (45) and PnB Designer (46), to confirm its on-target specificity (supplementary Table 8). Only one sgRNA (Figure 1C) appeared to be very specific for the W53X location as it would allow the binding of the spCas9 domain to the target locus that positions c.158G>A site within the editing window of ABE (4-8 for ABE8e, counting the PAM as 21-23). This sgRNA also had the highest on-target score (65.7) and lowest off-target score (56.8) among the Benchling-designed sgRNAs. The sgRNA targeting mouse *Kcnj13* was also selected based on the above criteria (highest on-target score; 57.1, lowest off-target score; 56.8). The chemically modified form of these sgRNAs (human; G*C*G*CUAGCGUUGGAUGAUGU and mouse; G*C*G*CUAGCGCUGGAUGAUGC) were ordered from Synthego (CA, USA).

### Generation of the lentiviral vector for Cas9-mediated gene editing

Lentivirus was manufactured for Cas9-mediated editing as described in Gándara, Carolina, et al. (2018). Briefly, the HEK293 cells were plated and ready for transfection after 24 h after plating. The target plasmid was transfected to the HEK293 cells at 70% confluence and packaging gene plasmids pMD2.G and psPAX2. The target vector plasmid contains a copy of the LCAguide, Cas9 protein, and mCherry reporter transgene driven by the EF-1α promoter. Cell culture supernatant was collected from HEK293 after transfection and was concentrated via gradient and centrifugation. A functional titer was performed to check the concentrated yield, between 10^7^ and 10^10^ particles/mL

### Gene editing in iPSC-RPE by lentiviral transduction (Cas9 and sgRNA delivery) and SNC (ssODN delivery)

For our attempt to edit the *KCNJ13* gene carrying a W53X nonsense mutation, we used viral transduction to deliver Cas9 and a sgRNA (TAATGGACATGCGCTAGCGT) to the mature iPSC-RPE cells. We used lentiviral vectors (100 MOI) designed explicitly for this purpose: lentiCRISPR v2-mCherry (Addgene plasmid # 99154), a gift from Agata Smogorzewska, with the annealed sgRNA oligonucleotides cloned into it using the BsmB1 enzyme digestion and ligation. Successful integration of the sgRNA sequence was confirmed by DNA sequencing with the primer 5′-GGACTATCATATGCTTACCG-3′ for the U6 promoter, which also drives sgRNA expression. Lentivirus was generated in-house using the method described above. This lentivirus was used to transduce mature iPSC-RPE cells to express Cas9, sgRNA, and the reporter gene mCherry, allowing easy identification of transduced cells. After 6 hours of viral transduction, 3 μg of the HDR repair template for W53X gene correction, ssODN-ATTO488 (GATGCTTGGGG**G**ATCCTAATGGA**T**ATGCGCT**G**GCGTTGGATGATGTT**A**GTCTTTTCTGCTTC T, the bold letters show the wobble changes), was delivered to the cells using SNC and incubated for 48 hours. Papain digestion was used to dissociate cells that expressed both red and green fluorescent markers, indicating that they had received both Cas9+sgRNA and ssODN constructs, into a single cell that was then used for patch-clamp experiments.

### Base editing in Kir7.1-HEK293 stable cells by electroporation

HEK293 stable cells expressing *KCNJ13^W53X^*-GFP were subcultured 24 hours before nucleofection at 70% confluency. The ABE8e mRNA (spCas9-NG, 3 µg) (47) and sgRNA (100 pmol) or ribonucleoprotein (RNP) complex formed by incubating the mixture of ABE8e protein (3 µg) and sgRNA (100 pmol) at room temperature for 10 mins were introduced via electroporation. 1×10^5^ cells were electroporated using the FS-100 program in the Lonza 4D nucleofector according to the manufacturer’s guidelines. Post-electroporation, cells were seeded in a 6-well plate and maintained in complete D-MEM media containing 100 µg/ml Hygromycin B for further analysis.

### Base editing in LCA16-patient’s specific fibroblasts and iPSC-RPE by SNCs

The W53X-LCA16-patient’s specific fibroblasts (Fibro^W53X^) were sub-cultured one day before treatment. For base editing, ABE8e mRNA (3 µg) and sgRNA (100 pmol) were delivered to fibroblasts using SNCs. 5 days post-treatment, DNA was isolated for genomic analysis. For base editing in iPSC-RPE, the cells were first seeded in a 96-well plate at a density of 50,000 cells per well in RDM containing 10% FBS and 10 µM ROCK inhibitor (R&D Systems; Cat# Y-27632). On Day 2, the media was switched to RDM containing 2% FBS. On day 3 post-seeding, ABE8e mRNA (3 µg) and sgRNA (100 pmol) were delivered to the cells using SNCs in RDM. the iPSC-RPE monolayer was dissociated 2 days post-treatment with SNC-ABE. Cells were seeded on transwell inserts and also collected for genomic DNA analysis. iPSC-RPE cells transitioned to transwell inserts and were cultured for 4-6 weeks to get a polarized monolayer of RPE and subsequently analyzed for Kir7.1 channel function by whole-cell patch-clamp approach. Untreated cells were used as references.

### Generating the Kcnj13^W53X/+^ Knock-in mouse model

The Kcnj13-W53X mouse model was generated by Cyagen Biosciences using CRISPR/Cas9-mediated genome engineering. Exon 2 was selected as the target site for the intended base change knock-in. Two distinct guides and a donor sequence. For homology-directed repair, the donor oligo carried the mutation p.W53* (TGG to TAG) flanked by 120 bp homologous sequences combined on both sides. Microinjections of Cas9 protein, sgRNA mixture, and ssODN were made into the pronucleus of fertilized eggs. The embryos were then transplanted to the pseudo-pregnant mice, and the resulting progeny were genotyped using PCR and RFLP to validate the desired gene mutation.

### Optical Coherence Tomography

Optical coherence tomography (OCT) was performed on mice anesthetized with ketamine/xylazine and whose pupils were dilated with 1% tropicamide using the Spectralis HRA+OCT system (Heidelberg Engineering Inc., Heidelberg, Germany). The mice were placed on a heating pad that was maintained at 37oC throughout the procedure, and a drop of artificial tear was applied before placing the corneal lens to keep the eyes from drying out. The images were captured and analyzed with the Heidelberg Eye Explorer software.

### Electroretinography (ERG) in mice

ERG was performed in mice using a standard protocol described elsewhere before and after the base editing to evaluate the function of the retina. Briefly, the mice were dark-adapted overnight prior to ERG. ERG signals were captured in full darkness using an Espion Ganzfeld full-field system (Diagnosys LLC, Lowell, MA). When using a contact electrode, a drop of 2% hypromellose solution was applied to the eye in order to keep the cornea wet and make electrical contact. Mice were subjected to multifocal electroretinogram (mfERG) testing with Celeris system (Diagnosys LLC, Lowell, MA), in which the retina is divided into 19 hexagonal areas and stimulated by a pseudo-random sequence of black and white hexagons that alternate multiple times per second. Data acquired were analyzed with Espion software (Diagnosys LLC, Lowell, Massachusetts) and Origin2018b OriginLab Corp., MA). For a and b-wave, the eyes were exposed to a series of flash intensities (0.03 to 30 cd.s/m2) using a ganzfeld dome for 400 ms with a 2 s interval between each flash. For c-wave, the eyes were exposed to 25 cd.s/m2 for 4 s. Animals were subjected to ERG every 2 weeks.

### Base editing in mice

C57BL/6J male and female mice (*Kcnj13^W53X/+^*) were housed at the animal facility at UW Madison (Madison, WI) under a 12-hour light-dark cycle at a controlled temperature (25±5 °C) and humidity (40– 50%). The mice were genotyped using standard PCR methods with the primers listed in supplementary Table 3, followed by digestion with restriction enzyme NheI (Anza# IVGN0066, ThermoFisher). The W53X mutation creates a restriction site for NheI, and therefore the W53X allele resulted in two (212 bp and 302 bp) fragments compared to only one (514 bp) fragment in the WT allele (Supplementary Figure 14). W53X-targeting sgRNA was designed and validated in mouse fibroblasts (48) isolated from *Kcnj13^W53X^*^/+^ mice. A subretinal injection (2 µl) of SNC-PEG-ATRA encapsulating ABE8e mRNA (2 and 3 µg), W53X-targeting sgRNA (100 pmol), and GFP mRNA (1 µg, to visualize the site of injection) was performed in mice (n=4 eyes). PBS or empty SNC-PEG-ATRA (n=4) injected eyes were used as reference. 5 days post-injection, imaging was carried out to assess the delivery based on GFP reporter mRNA. Genomic DNA was isolated from the optic cup of these mice to determine the on-target and off-target effects of base editing. To determine the editing efficiency in this monogenic W53X mouse model at the cellular level, any increase in the WT reads post base-editing of the W53X allele was noted and doubled to get the number of cells edited.

### On-target analysis by deep sequencing

Treated and untreated cells were dissociated using enzymatic treatment (Accutase/Papain) according to manufacturer’s instructions for genomic analysis. From the HEK293 stable cells, total RNA (Qiagen#74134) was isolated, and reverse transcribed to cDNA (ThermoFisher#4368814), and subsequently amplified for on-target analysis using *KCNJ13* Illumina-specific primers (supplementary Table 3). From fibroblasts, iPSC-RPE, and mouse optic cup, genomic DNA was isolated according to manufacturer’s guidelines (Quick-DNA^TM^ Miniprep plus kit #D4069) and quantified using Nanodrop 2000 or Qubit (Thermo Fisher). For deep sequencing of the *KCNJ13* locus, genomic DNA was amplified using Illumina-specific primers with adapter sequences (amplicon size ∽150 bp) (supplementary Table 4). Unique indexes (i7 and i5) were ligated to each custom amplicon by PCR (amplicon size 250 bp), and the indexed amplicons were pooled and purified using AMPure XP beads (Beckman Coulter #A63881). The indexed library was run on an Illumina MiniSeq instrument with a read length of 150bp. Deep sequencing data were analyzed using RGEN Cas-analyzer (49) (http://www.rgenome.net/cas-analyzer/) and CRISPResso2 (50) (https://crispresso.pinellolab.partners.org/submission) software.

### Off-target analysis by deep sequencing

The potential off-target sites for the hW53X-sgRNA were identified by an in-silico tool, Cas-OFFinder (36) (http://www.rgenome.net/cas-offinder/). The parameters used were an NG/NGG/NAG PAM with up to 4 mismatches to the sgRNA sequence (supplementary Table 5, Supplementary Figure 12). We also considered a DNA and RNA bulge of 1 nucleotide, which occurs due to an extra unpaired base in the DNA sequence concerning sgRNA or an extra unpaired base in sgRNA for DNA sequence in the genome, respectively. From the treated and untreated stable cells, fibroblasts, and iPSC-RPE, gDNA was isolated and amplified using primers specific to off-target sites. All the primer sequences are listed in supplementary Table 6. The deep sequencing and data analysis were performed as mentioned above.

### rhAmp off-target analysis in W53X mice

As the genomic DNA yield from the mouse optic cup was too low to amplify all the off-target sites separately, we used a highly efficient RNase H2-dependent (rhAmp) PCR technique that can amplify different targets in a single PCR reaction. Amplification and sequencing were performed according to ‘IDT’s rhAmp manual instructions). The rhAmpSeq CRISPR panel was designed using the IDT-designing tool (https://www.idtdna.com/pages/tools/rhampseq-design-tool) for the potential off-targets of mW53X-sgRNA identified using Cas-OFFinder (supplementary Table 7). The amplicon library was prepared using rhAmpSeq CRISPR Library Kit (IDT#10007317) and rhAmpSeq i5 and i7 index primers. The purified library was sequenced on the miniseq instrument from Illumina. Sequencing analysis was performed using IDT-rhAMP CRISPR analysis tool (https://www.idtdna.com/site/analysislab).

### Immunocytochemistry

Kir7.1 protein expression was assessed in the pool of W53X-mutant, WT, and base-edited HEK293 stable cells by immunocytochemistry as described earlier (10). As the protein is GFP tagged, GFP mouse monoclonal primary antibody (Cell Signaling#2955, 1:250) was used to enhance the Kir7.1 protein expression for its detection in the cells. Sodium Potassium ATPase rabbit monoclonal primary antibody (Thermo Fisher#ST0533, 1:500) was used to label the cell membranes. Alexa fluor-594 conjugated Donkey anti-Rabbit (Proteintech#SA00006.8, 1:500) and Alexa fluor-488 conjugated Donkey anti-Mouse (Proteintech#SA00006.5, 1:500) secondary antibodies were used. DAPI was used as a nuclear counterstain. Immunostained cells were imaged on a confocal microscope (Nikon C2 Instrument).

### Electrophysiology assay

A high throughput automated patch clamp (Q Patch II, Sophion, Denmark) was used to measure the whole-cell current from the HEK^WT^, HEK^W53X,^ and base-edited HEK^W53X^ stable cells as described earlier (51). Briefly, the cells were grown in a T75 flask for 48-72 hours and then detached gently using Detachin^TM^. The cells were centrifuged at 90 x g for 1 min and resuspended in serum-free media containing 25 mM HEPES. The cells (3 M/ml) were kept on a shaker for 20 mins before the experiment. 48 cells were recorded in parallel on single-hole disposable Qplates with individual amplifiers. A pressure protocol was used to achieve cell positioning (−70 mbar), Giga seal (−75 mbar), and whole-cell configuration (5 pulses with −50 mbar increment between the pulses, first pulse of −250 mbar). The current was recorded in response to voltage-clamp steps from the holding potential (−10mV) to voltages between −140mV and +40mV (Δ=10mV). More than 70% of the cells completed the experiment. The cells in which the stability was compromised during the experiment were judged by the leak current and excluded from the analysis. The extracellular solution contained (in mM): 135 NaCl, 5 KCl, 10 HEPES, 10 glucose, 1.8 CaCl_2,_ and 1 MgCl_2,_ pH adjusted to 7.4 and osmolarity 305 mOsm. The intracellular solution contained (in mM) 30 KCl, 83 K-gluconate, 10 HEPES, 5.5 EGTA, 0.5 CaCl_2_, 4 Mg-ATP, and 0.1 GTP, pH adjusted to 7.2 and osmolarity 280 mOsm. In an alternative external solution, NaCl was replaced with RbCl (140 mM) and used as an enhancer of Kir7.1 current. An extracellular solution with 20 mM Cs^+^ was used to block the Kir7.1 current. The data was analyzed using Sophion Analyzer v6.6.44. Whole-cell manual patch-clamp recording of base-edited hiPS-RPE cells was performed according to the standard protocol described elsewhere (7). There was no reporter or selection marker to aid in identifying the recipient or edited cells. Therefore, these cells were picked up randomly for the electrophysiology assay.

### Statistical analysis

The data analysis was done using Origin software (Origin 2020, OriginLab Corporation, Northampton, MA, USA) and expressed as mean ± standard error. Two-tailed unpaired Student’s *t*-test was used to determine the statistical differences, and *p* < 0.05 was considered significant.

### Study approval

All work with LCA16 patient-derived cells (fibroblasts, iPSCs, and iPSC-RPE) was carried out following institutional, national, and international guidelines and approved by the University of Wisconsin-Madison’s institutional review board and Stem Cell Research Oversight Committee. The animal protocols followed the ARVO Statement for use in ophthalmic and vision science research and were approved by the University of Wisconsin School of Medicine and Public Health Animal Care and Use Committee (ACUC).

## Data availability

Data is available in the paper’s supplemental material, and additional data can be made available from the corresponding author upon request.

## Author contributions

M.K., and P.K.S. drafted the manuscript.

M.K, P.K.S, Y.W., D.S., G.A.N., A.A.A., D.M.G., D.R.L., S.G., K.S., and B.R.P. conceptualized the study.

M.K, P.K.S, Y.W., D.S., G.A.N., A.A.A., D.M.G., D.R.L., S.G., K.S., and B.R.P. devised the methodology

M.K., P.K.S., Y.W., D.S., A.S., G.A.N., S.S., Y.T., Y.C., and A.A.A. carried out the investigation.

M.K, P.K.S., D.M.G., S.G., K.S., and B.R.P. validated the study.

M.K, P.K.S, and Y.W. carried out the formal data analysis.

The data was visualized by M.K, P.K.S., Y.W., D.M.G., S.G., K.S., and B.R.P..

D.S., G.A.N., Y.T., K.L.E., and C.O.T., and D.R.L. provided the resources.

M.K., P.K.S., Y.W., G.A.N., D.M.G., D.R.L., S.G., K.S., and B.R.P. curated the data.

D.M.G., D.R.L., S.G., K.S., and B.R.P. supervised the study.

D.M.G., D.R.L., S.G., K.S., and B.R.P. acquired the funding.

## Acknowledgments

We thank the Stem Cell & Regenerative Medicine Center (SCRMC) fellowship for the funding support to Meha Kabra. We acknowledge Retina Research Foundations Kathryn and Latimer Murfee Chair (KS), and M.D. Matthews Research Professorship (BRP), Retina Research Foundation Emmett A. Humble Distinguished Directorship (DMG), and McPherson ERI Sandra Lemke Trout Chair in Eye Research (DMG). The research was also supported by UW-Partnership Education and Research Committee funds and Harrington Discovery ‘Institute’s Gund-Harrington Scholar (KS). We thank the Helen Hay Whitney Foundation, HHMI, and the K99 award HL163805 for support to GAN. This work was supported by NIH awards NIH R01 EY024995 and R24 EY032434 (BRP), U01 AI142756, RM1 HG009490, and R35 GM118062 (DRL), and Research to Prevent Blindness. This study was partly supported by S10OD018221 and the Core Grant (P30 EY016665) for Vision Research from the NIH to the University of Wisconsin-Madison.

## Supplementary Note

### Inefficient repair of W53X in patient iPSC-RPE^W53X^ using CRISPR-Cas9 nuclease

We first assessed the correction of the homozygous *KCNJ13* W53X mutation in iPSC-derived RPE (iPSC-RPE^W53X/W53X^) using Cas9 nuclease-mediated HDR. We delivered Cas9 nuclease and sgRNA via the lentiviral vector, LentiCRISPRv2-mCherry (Supplementary Figure 1), and the HDR donor template (ssODN-ATTO488) via SNC. The size and zeta-potential of donor templated-encapsulated SNC were summarized in supplementary Table 1. We confirmed the delivery of both constructs using their cognate fluorescent reporters (Supplementary Figure 2A). Deep sequencing analysis of the treated samples indicated that most of the indels (5.05 ± 1.24%) were created downstream of the pathogenic mutation (TAG), resulting in no change to the reading frame (Supplementary Figure 2, B and C). Only a small fraction of reads showed in-frame indel formation (1.03 ± 0.52%) and a corrected wildtype (WT) genotype (0.34%) (Supplementary Figure 2D), both of which are predicted to remove (i.e., repair) the W53X stop codon during translation of the edited *KCNJ13* mutant allele. Although the green fluorescence in the treated cells showed a successful delivery of ssODN, we did not observe the inclusion of any silent nucleotide bases from the ssODN (Supplementary Figure 2B). Most reads (94.50 ± 1.25%) were unedited in the treated cells.

Next, we measured the Kir7.1 channel function in gene-edited cells expressing the red (mCherry) and green (ATTO488) fluorescence, indicating that they had received both Cas9-sgRNA and ssODN. Single-cell patch-clamp recordings revealed a normal Kir7.1 current with an inward current of −101.1 ± 35.54 pA at −150 mV in treated cells. Substitution of extracellular Na^+^ with rubidium (Rb^+^), a known activator of Kir7.1, increased the Kir7.1 inward current by 7-fold to −713.5 ± 92.97 pA (Supplementary Figure 2, E-G). These results indicate that delivery of both Cas9-sgRNA and ssODN can sometimes generate function-restoring gene edits. However, the exact nature and frequency of editing outcomes in the single cells we recorded by patch clamp could not be assayed due to the technical challenges of amplifying genomic DNA from a single cell.

Altogether, this nuclease-mediated HDR approach did result in a very small frequency of functional RPE cells– indicating that gene correction is a viable strategy to rescue Kir7.1 function in RPE cells. However, the low efficiency and high heterogeneity of the edits diminished the translational potential of this strategy.

## Supplementary tables

**supplementary Table 1:** Size and zeta-potential of ATRA-modified SNCs with different.

| <u>Payload</u> | <u>Size ($D_h$, nm)</u> | <u>Zeta-potential (mV)</u> |
| --- | --- | --- |
| <u>ssODN donor template</u> | <u>46±4</u> | <u>3.6±1.6</u> |
| <u>ABE mRNA+sgRNA</u> | <u>42±4</u> | <u>4.9±1.8</u> |

**supplementary Table 2:**
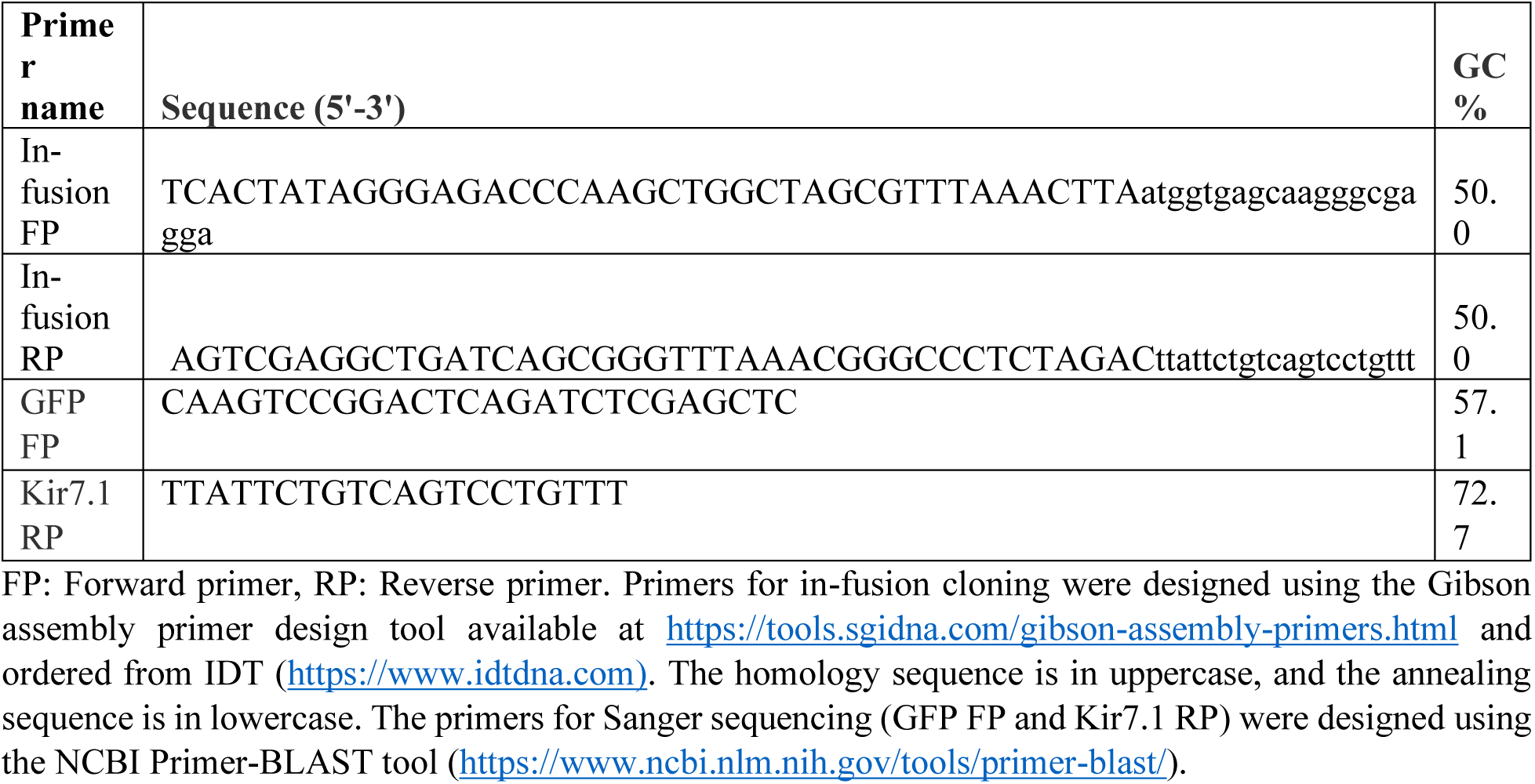
Primers for in-fusion cloning of *KCNJ13* in FLP-In™ expression vector.

**supplementary Table 3:**
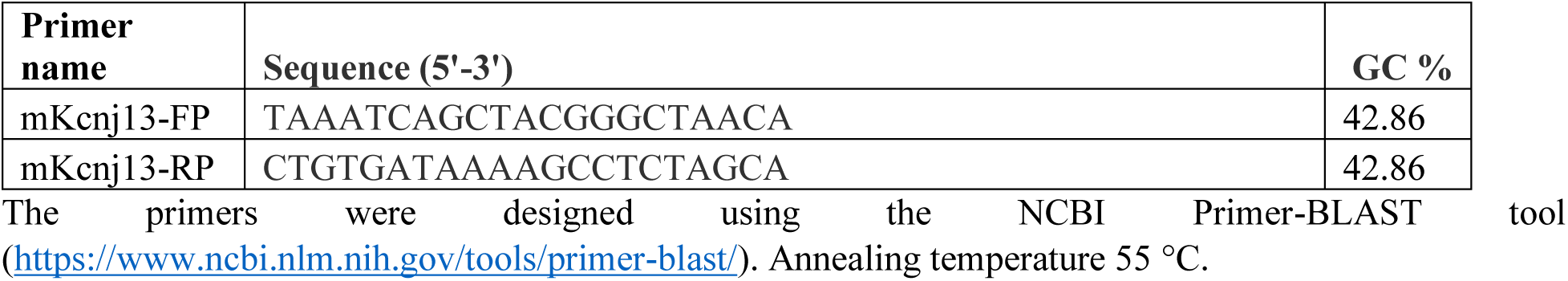
Primers to genotype mice.

**supplementary Table 4:**
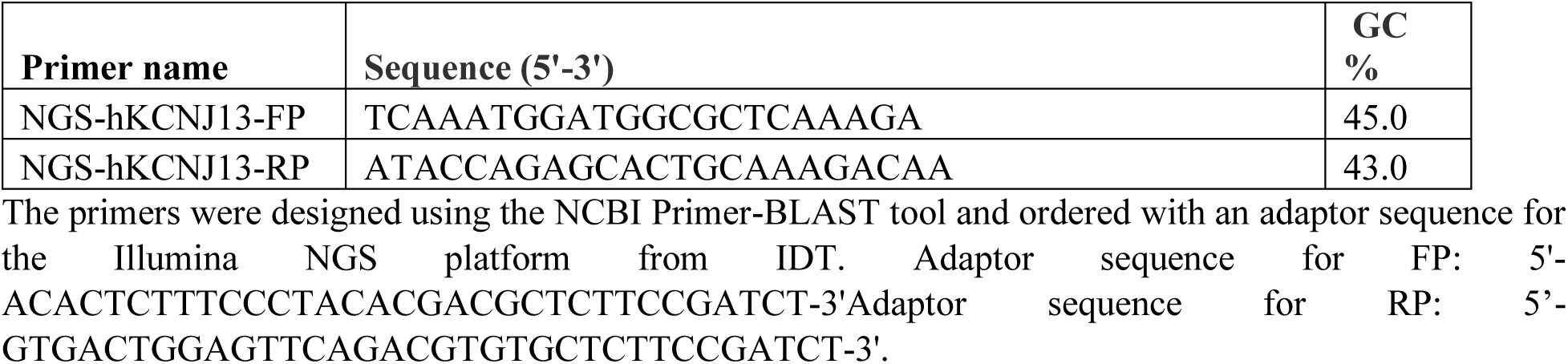
Primers to amplify the *hKCNJ13* on-target sites.

**supplementary Table 5:**
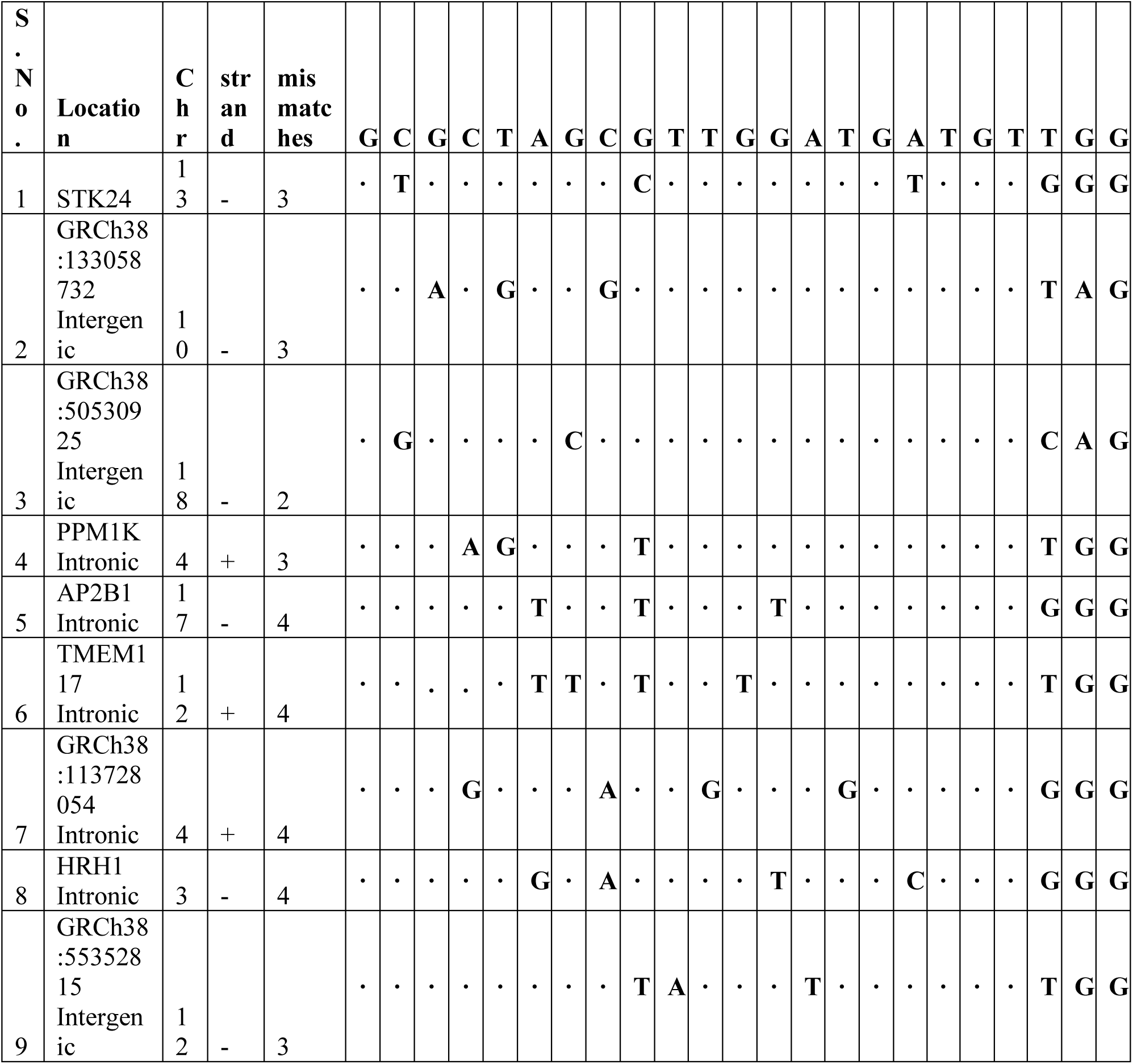
Putative off-targets sites of human-sgRNA identified using Cas-OFFinder.

**supplementary Table 6:**
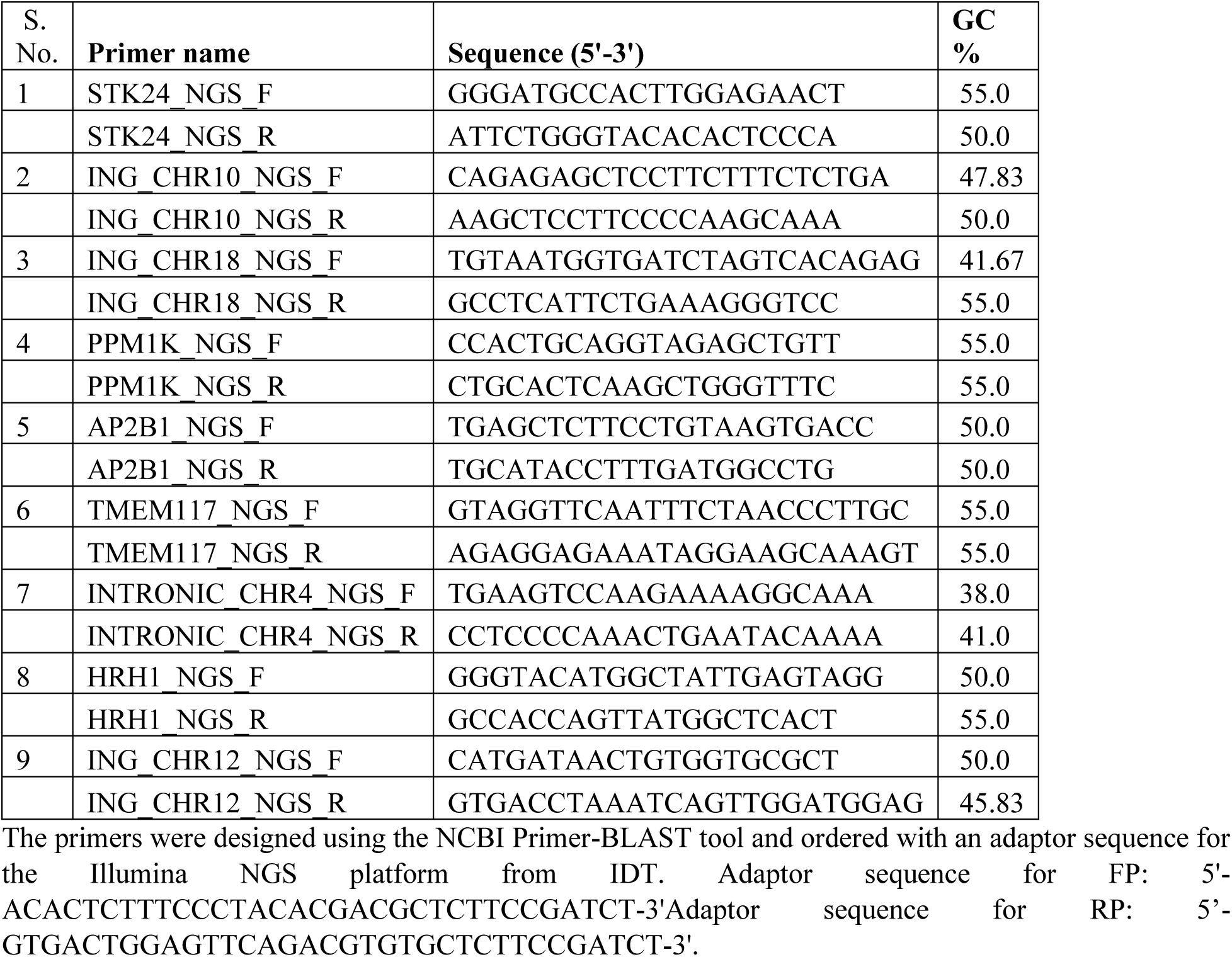
Primers to amplify the off-target sites of human-sgRNA.

**supplementary Table 7:**
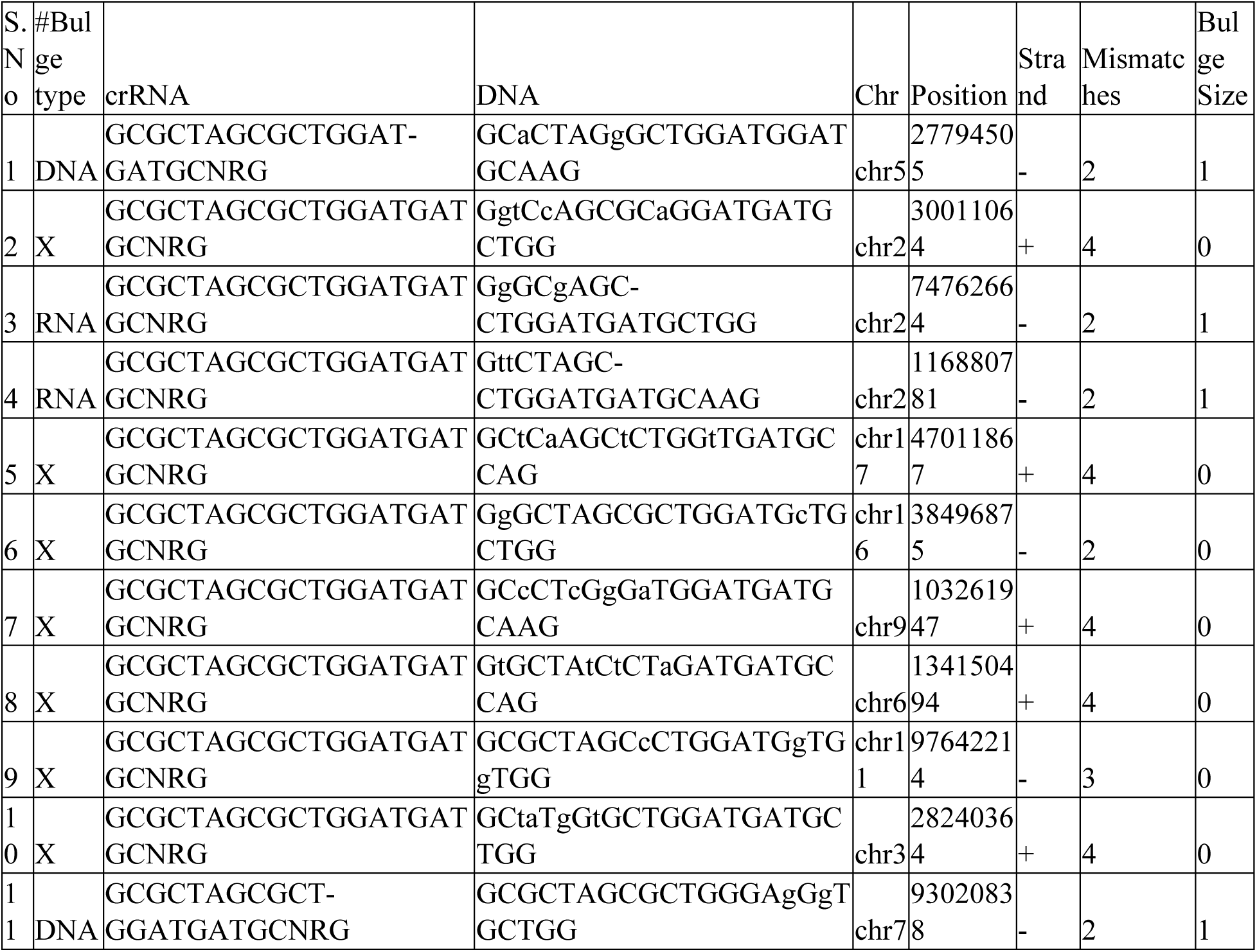
Potential off-target sites of mouse W53X-sgRNA identified using Cas-OFFinder.

**supplementary Table 8:**
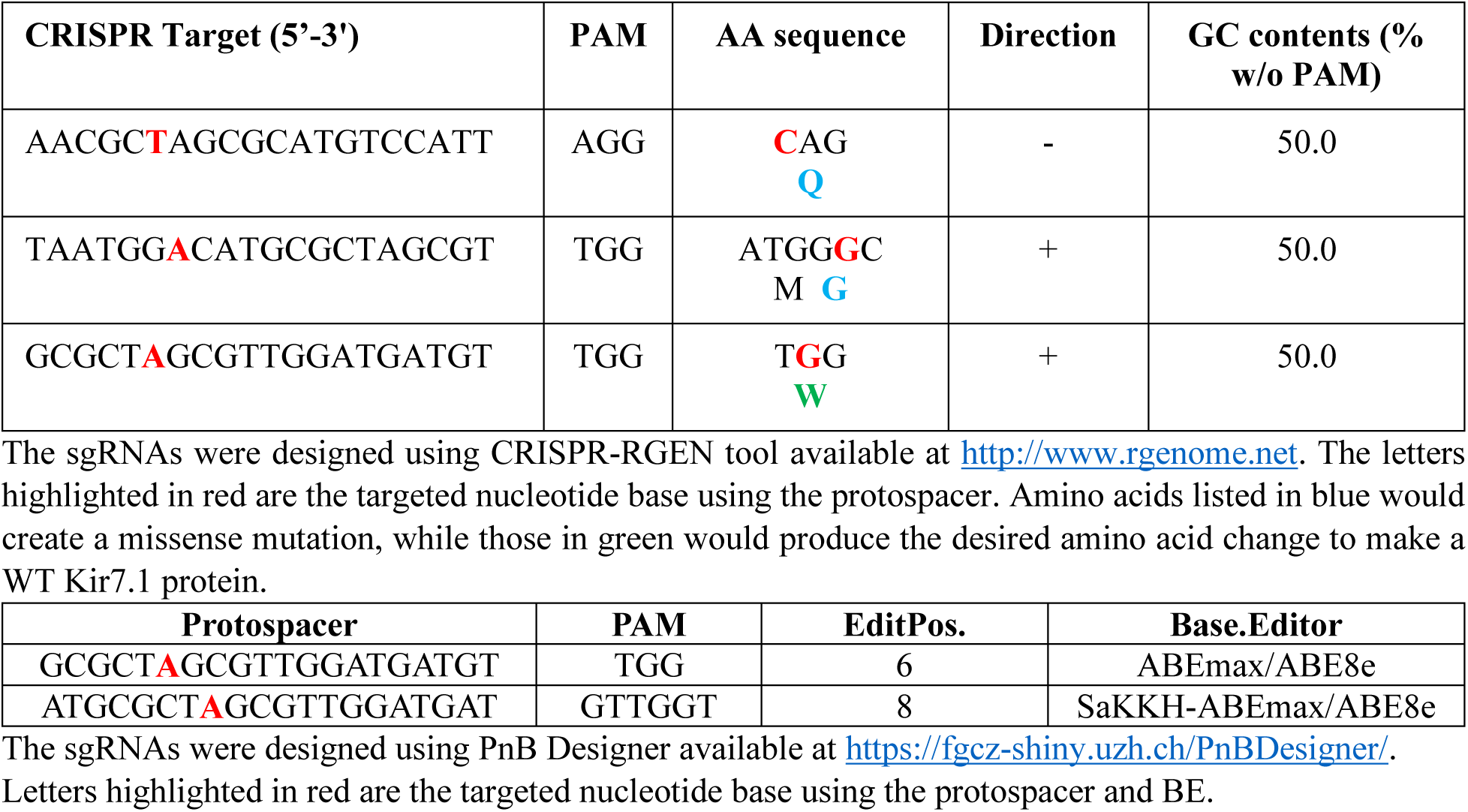
sgRNA design using CRISPR-RGEN and PnB Designer.

**supplementary Table 9:**
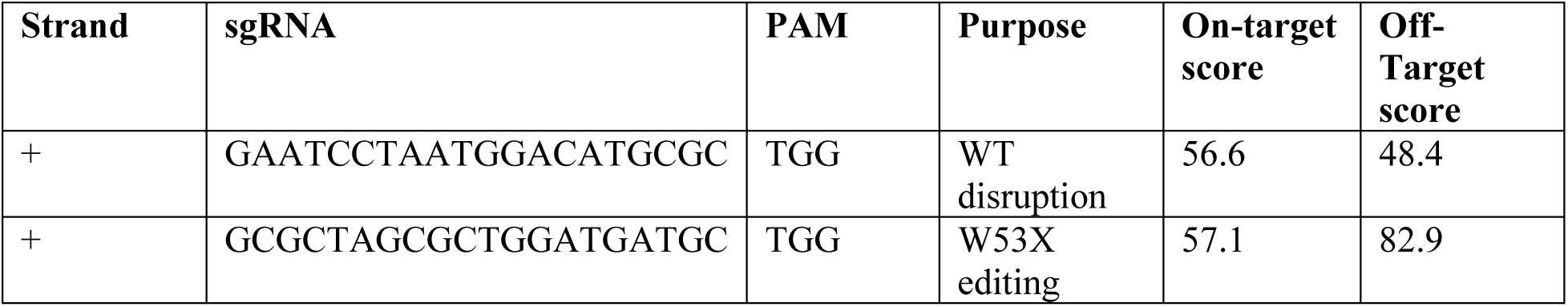
sgRNA design for mouse *Kcnj13* using Benchling.

## Supplementary Figures

**Supplementary Figure 1:**
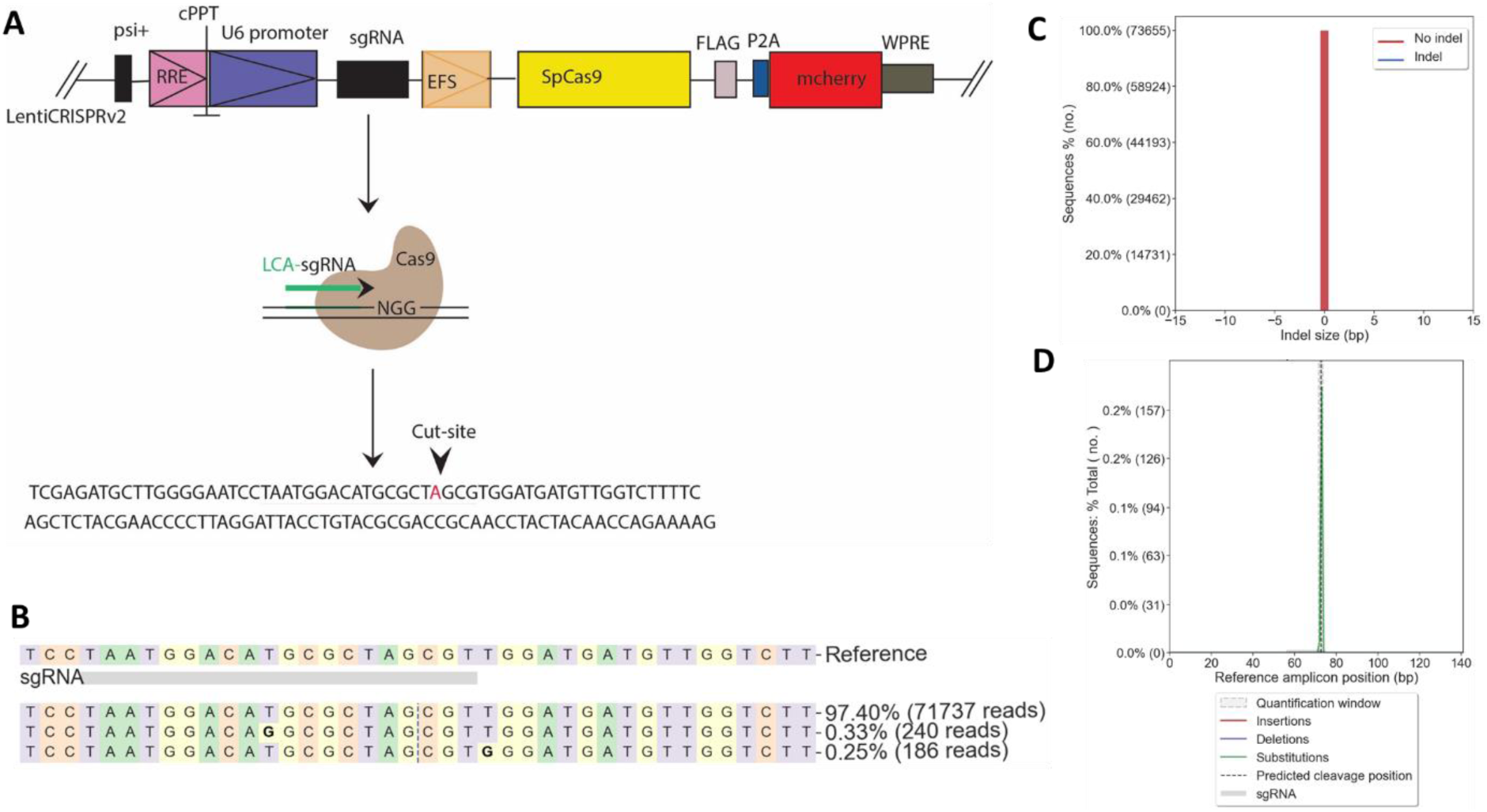
Gene editing strategy and sequencing readouts from untreated iPSC RPE^W53X^. A] Construct design of lentiviral CRISPRv2 vector demonstrating the sgRNA and spCas9 location. B] Nucleotide distribution around the cut site (vertical dashed line) of sgRNA. C] Indel size distribution in untreated samples. D] Mutation position distribution showing the editing profile in untreated samples.

**Supplementary Figure 2:**
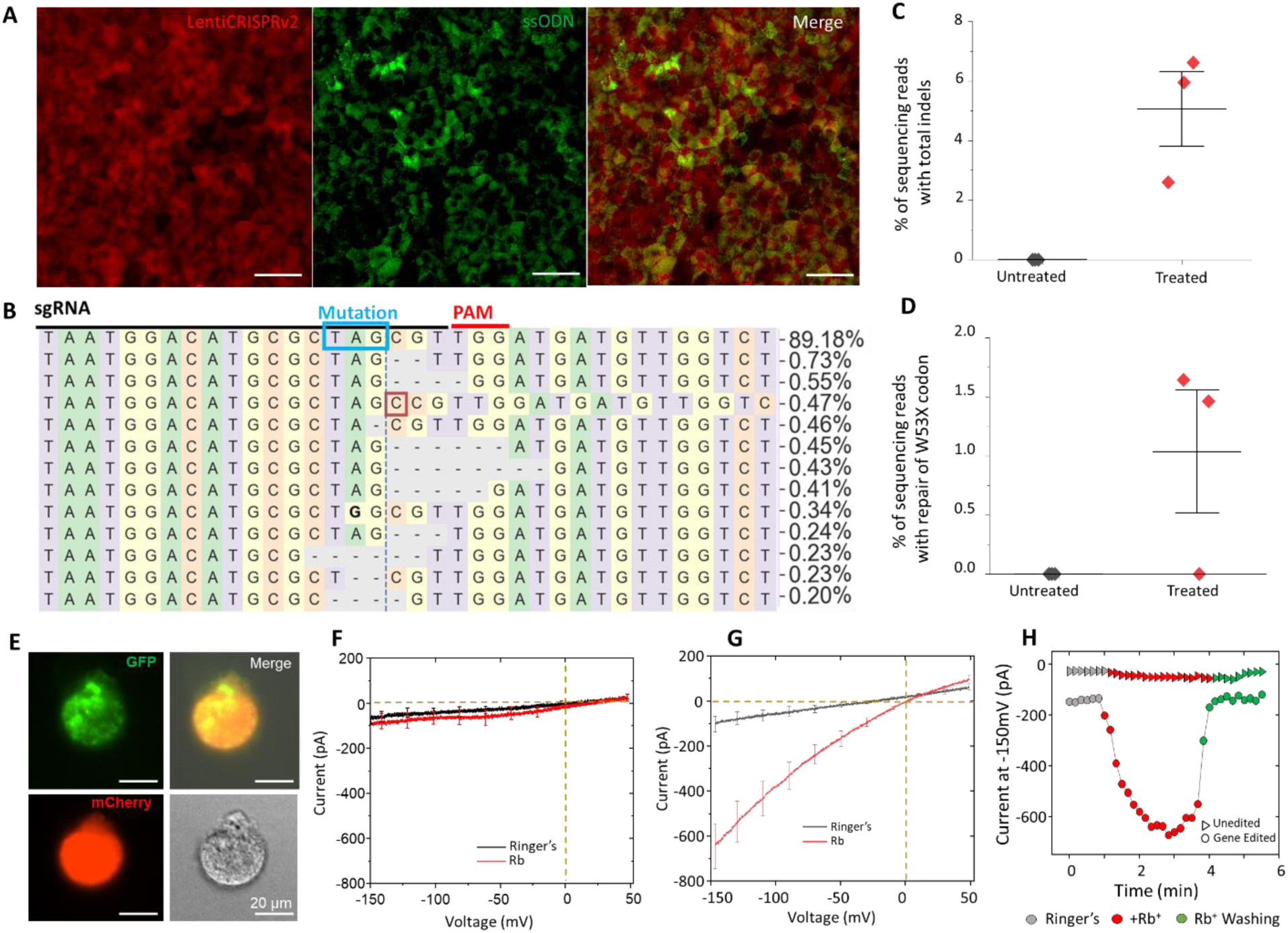
CRISPR-Cas9 nuclease editing outcomes in iPSC-RPE^W53X/W53X^. **(A)** Transduction with LentiCRISPRv2 –mCherry carrying Cas9 and sgRNA (red panel) and delivery of ssODN-ATTO488 via SNCs (green panel) in iPSC RPE^W53X/W53X^ showing the dual fluorescence (merged panel). Scale 50 µm. (**B)** Deep sequencing reads from cells that received both LentiCRISPRv2 and ssODN treatment show the nucleotide distribution around the DSB cleavage site (vertical dashed line). The sgRNA protospacer location is highlighted by a black line, the protospacer adjacent motif (PAM) by a red line, and the pathogenic early stop codon (TAG) mutation in the blue box. Substitutions are highlighted in bold, insertions are shown in red boxes, and a dash shows deletions. (**C)** Total percentage of sequencing reads containing indels at the DSB site across the treated samples (n=3, biological replicates). (**D)** Percentage of reads comprised of WT reads and in-frame indels observed in the treated samples (n=3, biological replicates). **(E)** Fluorescence microscopy image of a single dissociated iPSC-RPE cell expressing both reporters, chosen for manual patch-clamping. Scale 20 µm. **(F)** Average current-voltage plot from untreated cells used as a reference. The black line indicates the current recorded in HEPES ‘Ringer’s (HR) solution and red line indicates the current recorded in the presence of extracellular Rb^+^ ion. (n=3). **(G)** Average current-voltage plot demonstrating K^+^ current following gene editing. The grey line indicates the current recorded in the HR solution, while the red line indicates the current measurement in the presence of the Kir7.1 channel current enhancer, Rb^+^ ion (n=4). **(H)** A representative time course illustrating the reversible increase in inward current by extracellular (HR, grey), Rb^+^(red), and wash (green).

**Supplementary Figure 3:**
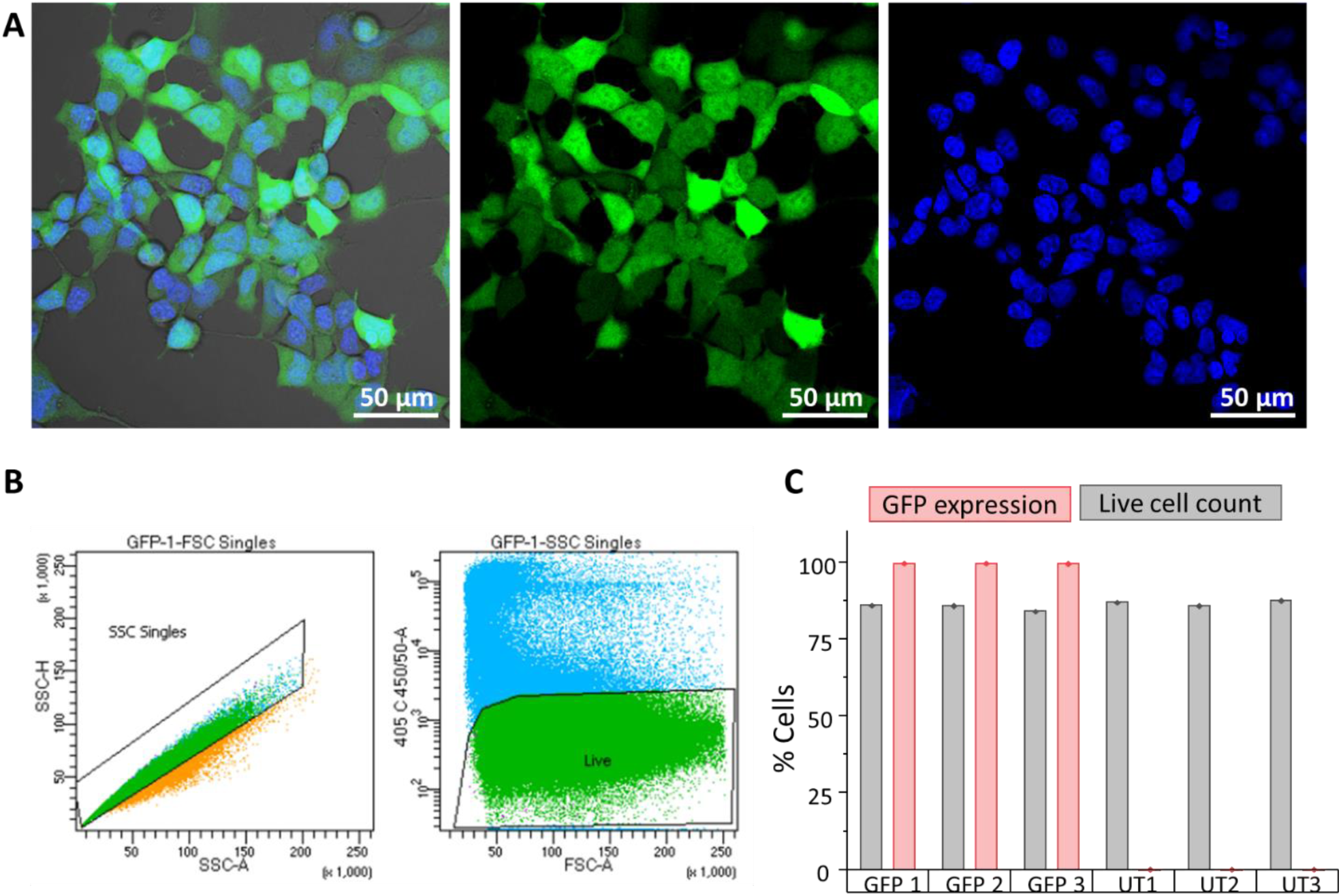
Electroporation efficiency in HEK293 cells. A] Confocal imaging of HEK293 cells electroporated with GFP mRNA showing the efficient delivery of GFP (green). Hoechst stain (blue) was used to label nuclei. B] Fluorescence-activated cell sorting (FACS) of GFP-electroporated cell population. Single cell population by forward scatter (FSC) and side scatter (SSC). C] % of live and GFP positive cells from a cell population either electroporated with GFP (n=3) or placebo treated (n=3).

**Supplementary Figure 4:**
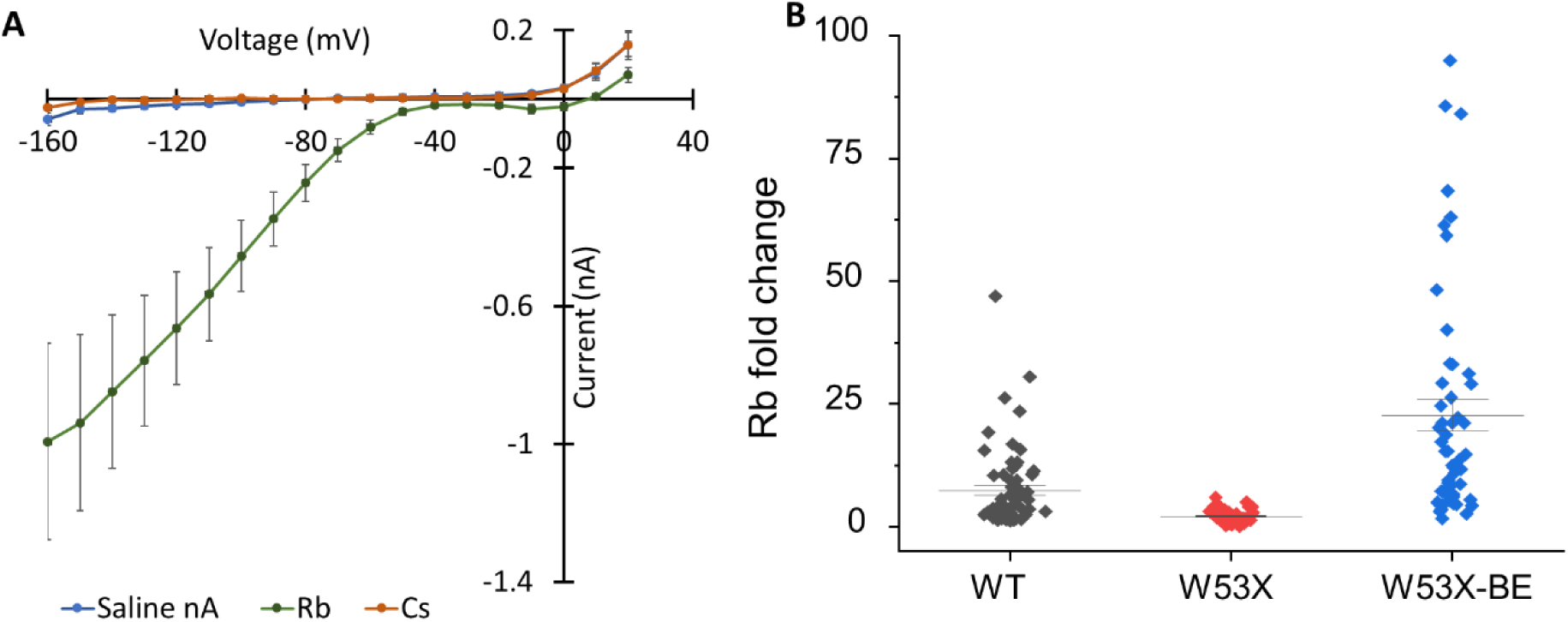
Automated patch-clamp recordings from HEK^W53X/W53X^, HEKT^WT/WT,^ and base-edited HEK^W53X^ cells. A] I-V curve for HEK^WT/WT^ cells showing a large negative membrane potential. B] The fold changes for Rb^+^ in HEK^WT/WT^, HEK^W53X/W53X^, and base-edited HEK^W53X^ cells.

**Supplementary Figure 5:**
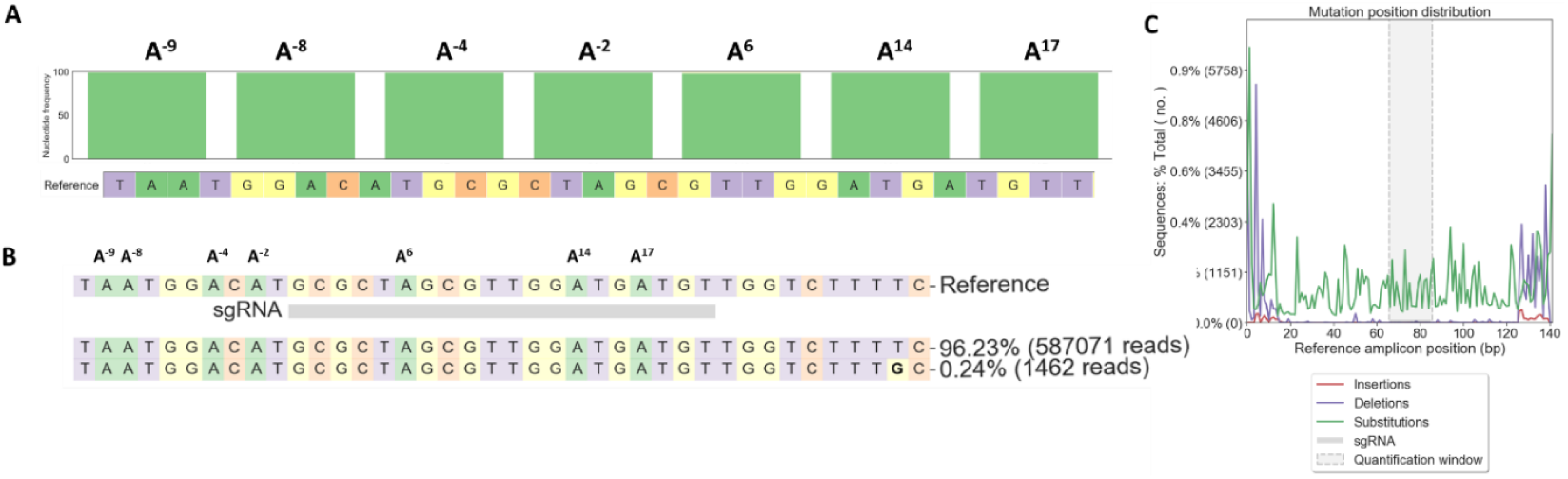
Sequencing readouts from untreated LCA16-Fibroblasts^W53X^ used as reference. A] Nucleotide distribution around sgRNA location as observed in sequencing reads. B] Percentage of sequencing reads observed in the untreated sample. C] Percentage distribution of substitution and deletion at sgRNA location.

**Supplementary Figure 6:**
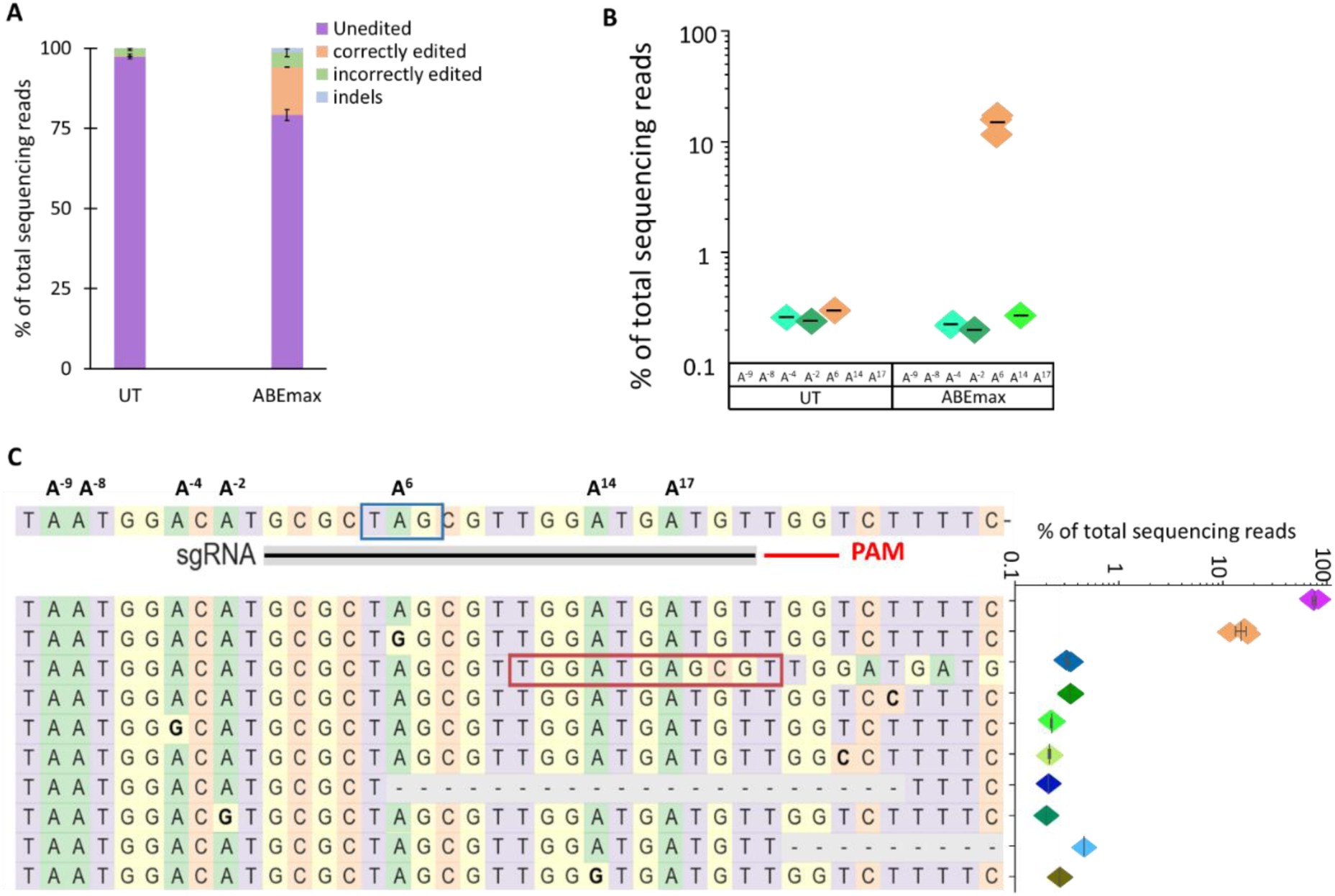
Evaluation of ABE7.10max mRNA + sgRNA combinations to correct the W53X allele in Fibro^W53X^. A] Base editing efficiencies shown as the % of total DNA sequencing reads, classified as unedited, correctly edited, incorrectly edited due to bystander ‘A’ edits, and with indels in treated and untreated (UT) cells. B] % Editing of the target (A^6^) and bystander (A^-9^, -A^-8^, A^-4^, A^-2^, A^14^, A^17^) ‘A’ to ‘G’ as observed in three independent experiments. C] The sgRNA location is marked by black line, PAM by red line, and mutation in the blue box. All the ‘A’ bases within the protospacer are numbered from 1-20 based on their location. The ‘A’ bases downstream of the protospacer are numbered from −1 to −9, considering +1 as the first base of the protospacer. The top 10 most frequent alleles generated by ABE7.10max mRNA treatment show the nucleotide distribution around the cleavage site for sgRNA. Substitutions are highlighted in bold, insertions are shown in the red box, and deletions are shown by dashes. The scatter plot shows the frequency of reads observed in treated cells (n=3 biological replicates). Figures presenting data from replicates are shown as mean ± SEM.

**Supplementary Figure 7:**
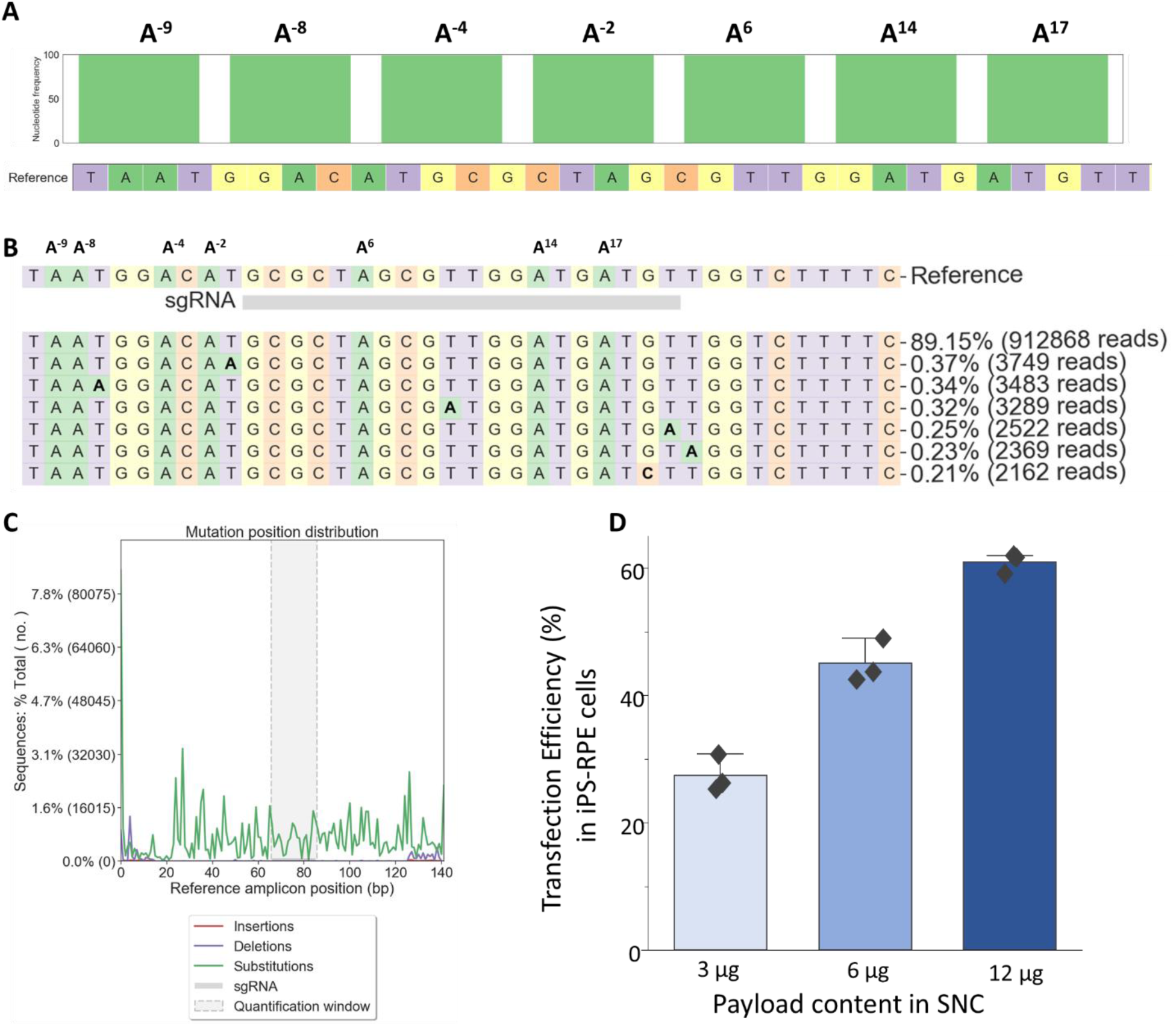
Sequencing readouts from untreated iPSC RPE^W53X^ cells used as reference. A] Nucleotide distribution around sgRNA location as observed in sequencing reads. B] Percentage of sequencing reads observed in the untreated sample. C] Percentage distribution of substitution and deletion at sgRNA location. D] Transfection efficiency in iPSC-RPE cells using SNCs.

**Supplementary Figure 8:**
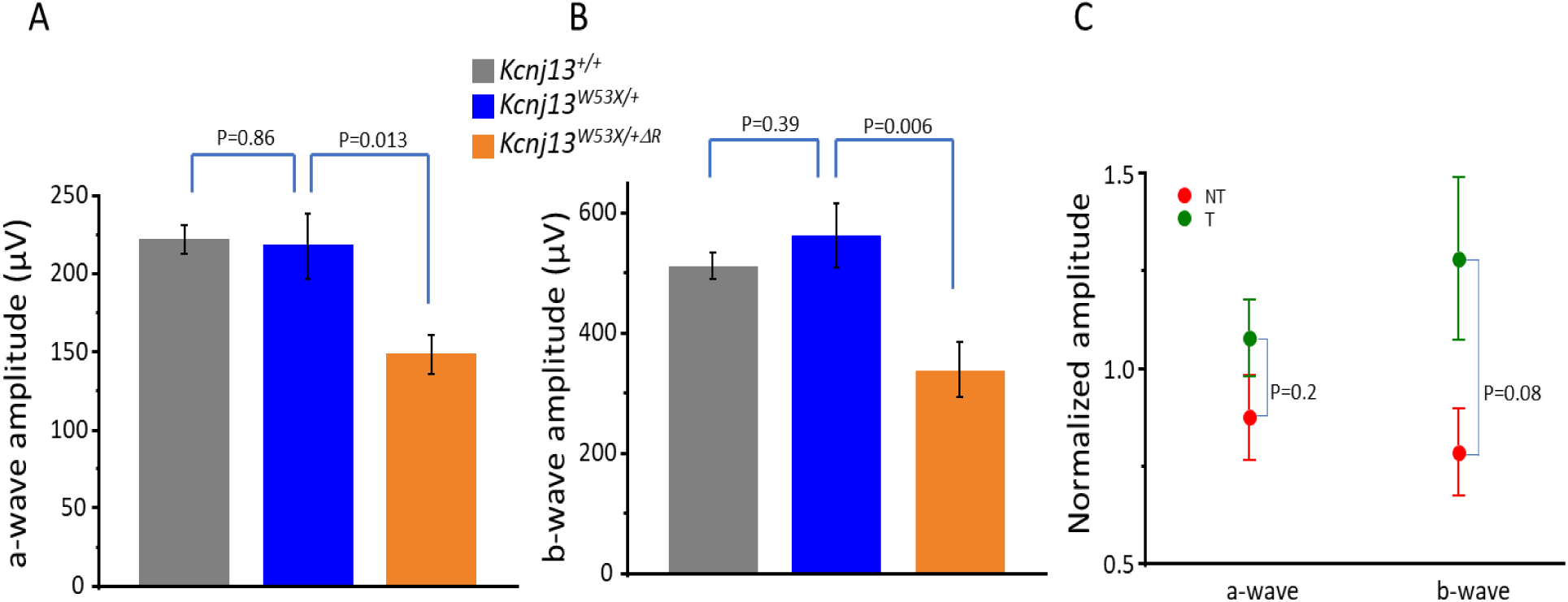
Comparison of a-wave and b-wave amplitude: A] a-wave B] b-wave amplitude comparison after scotopic ERG on Kcnj13^+/+^, Kcnj13^W53X/+^ and Kcnj13^W53X/+ΔR^ mice. C] comparison of a-wave and b-wave amplitudes in Kcnj13^W53X/+ΔR^ following the injection of base editor.

**Supplementary Figure 9:**
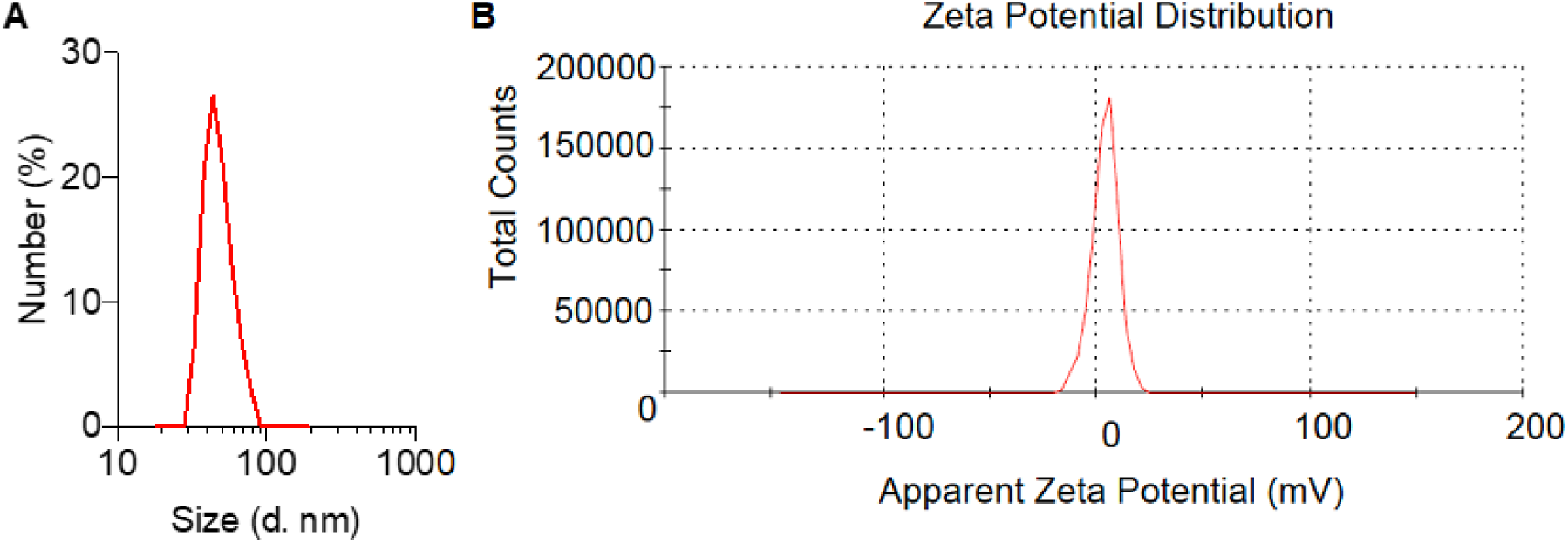
**A]** Size distribution and **B]** Zeta-potential of ABE mRNA+sgRNA-encapsulated SNC with ATRA modification.

**Supplementary Figure 10:**
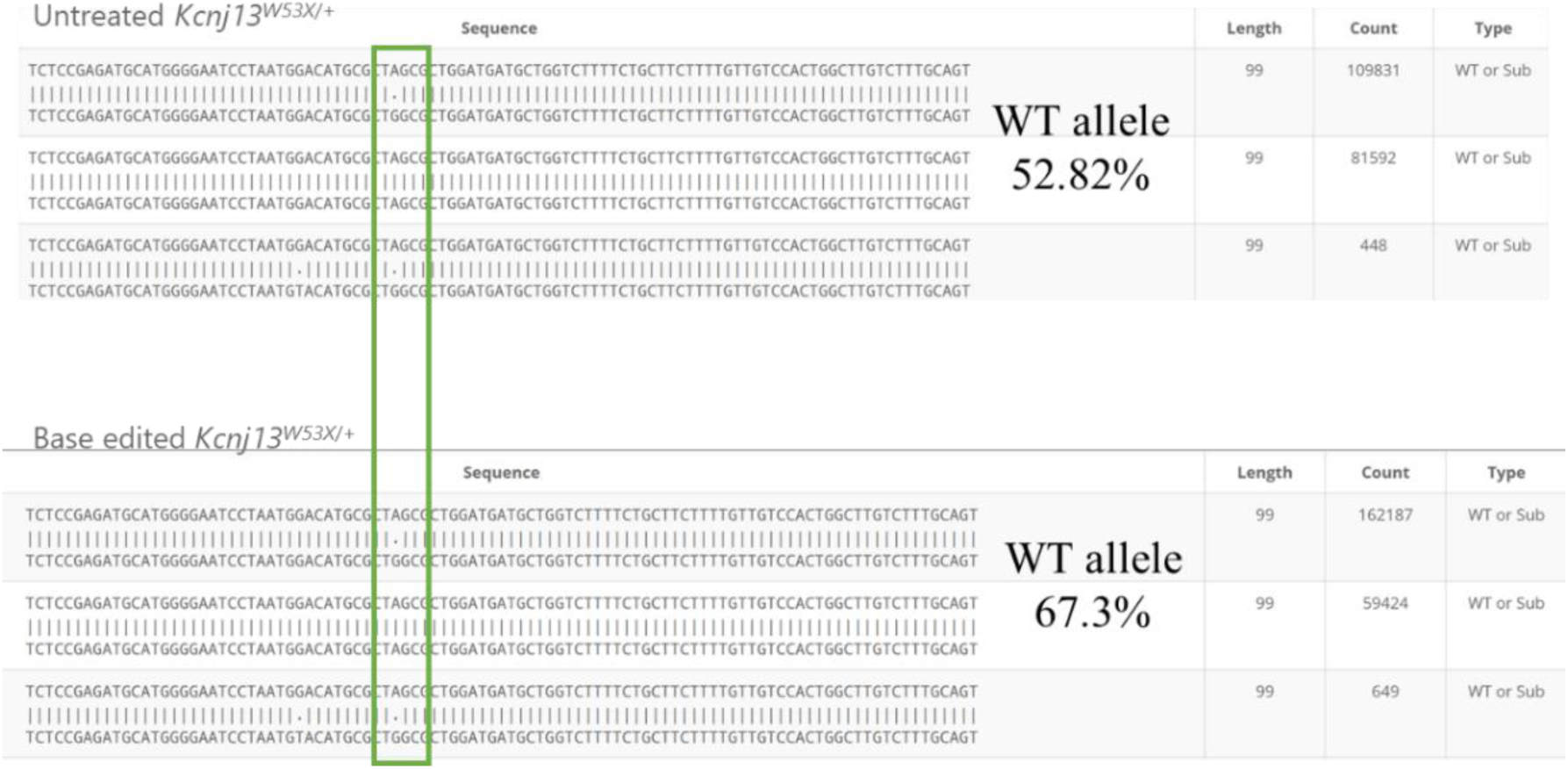
Deep sequencing reads from the edited fibroblasts isolated from Kcnj13^W53X/-^ mice. The sequencing reads were generated by editing mouse W53X alleles within fibroblasts using ABE8e and W53X-sgRNA, delivered via nucleofection.

**Supplementary Figure 11:**
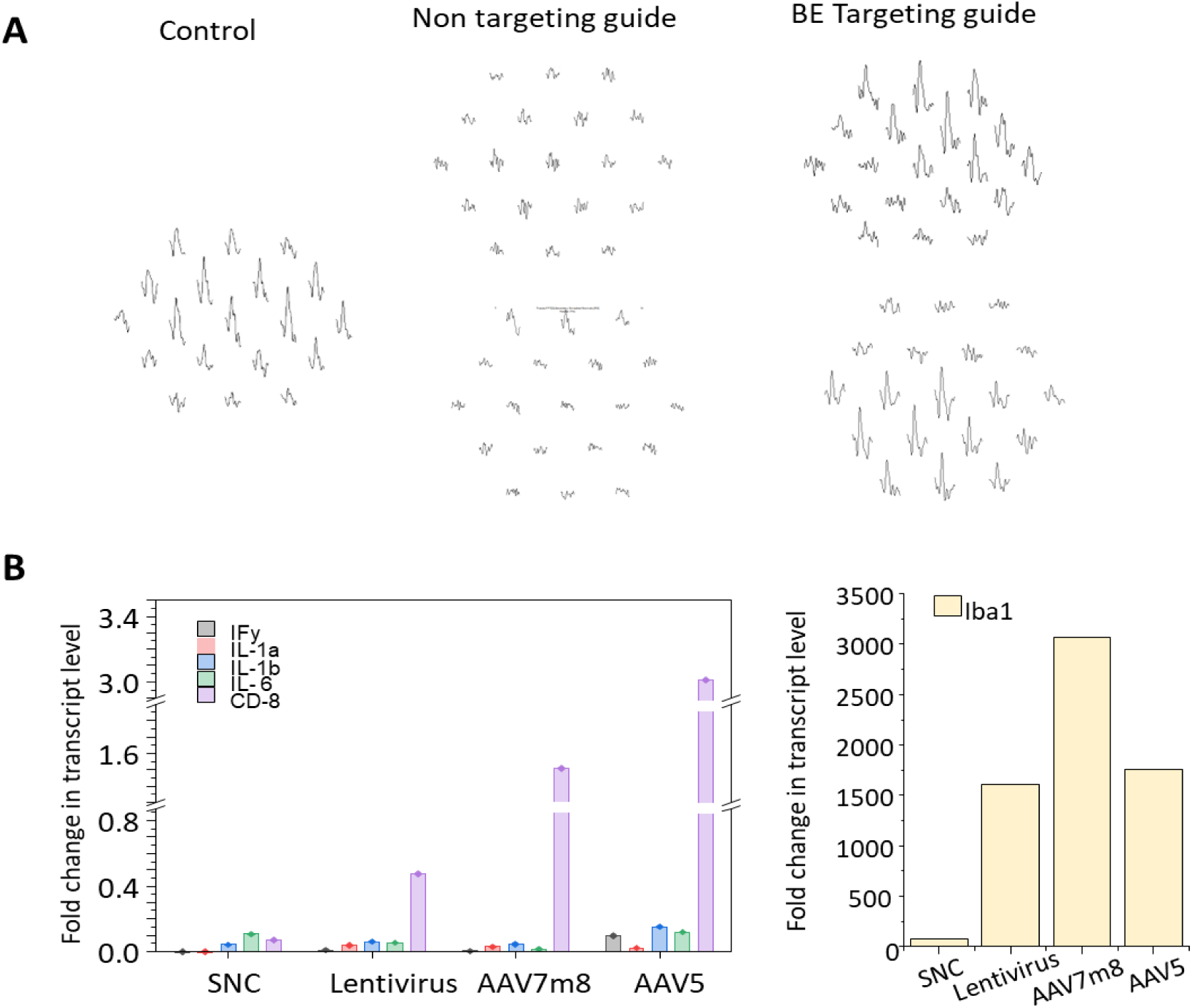
A] mfERG measurements from the control mice (left panel), after wildtype allele disruption (middle panel) and after injection of base editor to Kcnj13^W53X/+ΔR^ mice (right panel). B] Level of inflammatory cytokine transcripts triggered by SNC, Lentivirus, AAV7m8, and AAV5.

**Supplementary Figure 12:**
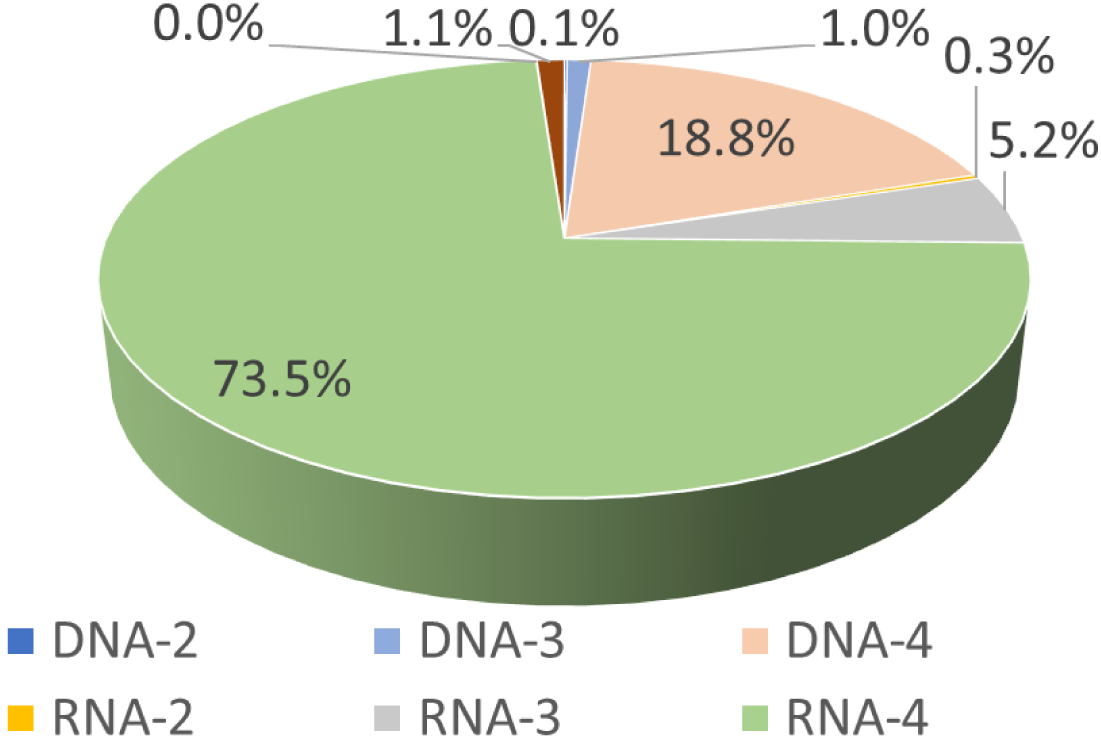
Genomic off-target sites for human W53X sgRNA. The sites were identified using standard criteria, up to 1-4 mismatches and DNA/ RNA bulge (size=1 nucleotide).

**Supplementary Figure 13:**
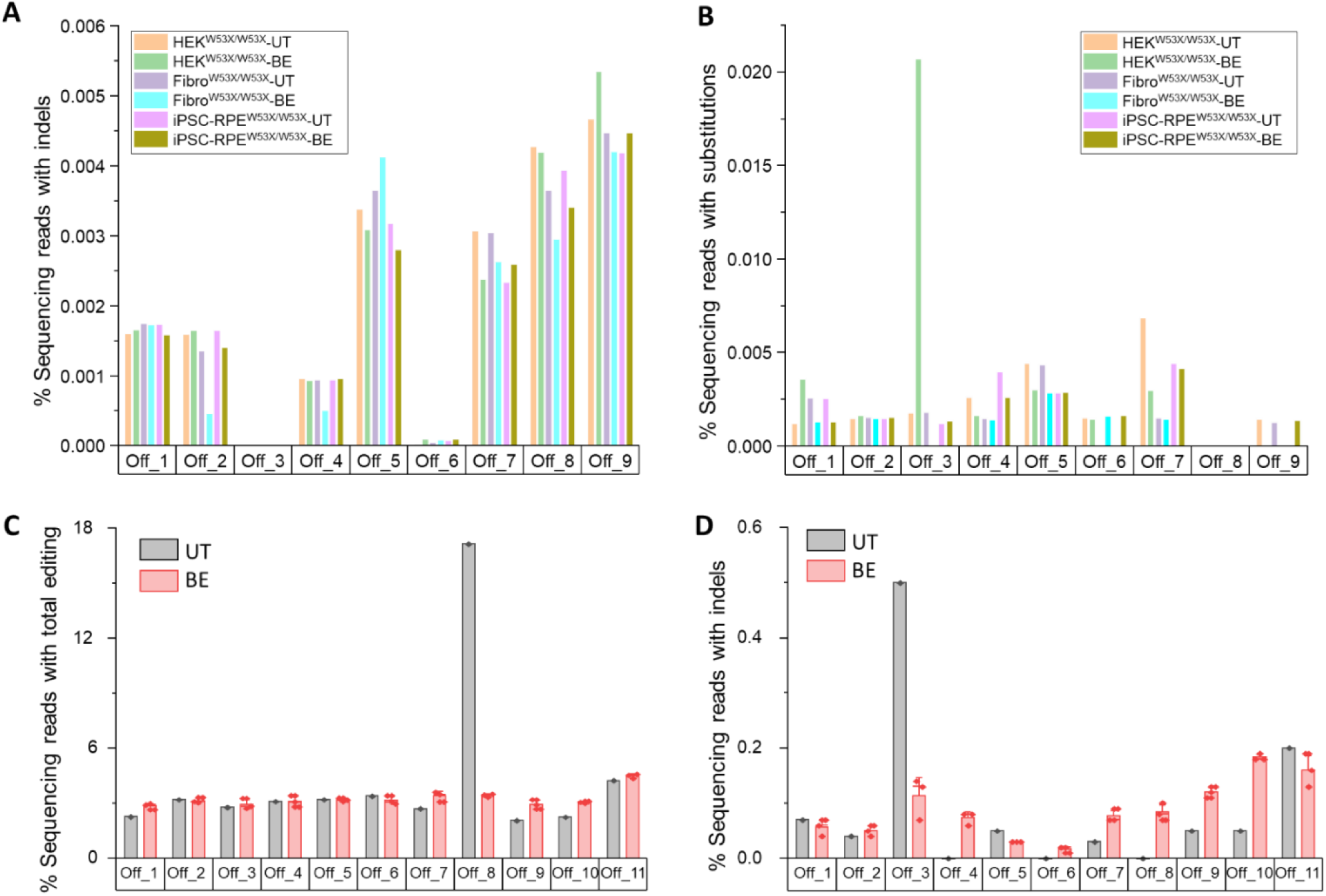
Off-target editing analysis of the base editor delivered by SNC in the human in vitro LCA16 model and mouse in vivo LCA16 model. **(A)** Percentage of sequencing reads with indels as observed by deep sequencing analysis in Base editor treated (BE) HEK^W53X^, Fibro^W53X,^ and iPSC-RPE^W53X/W53X^ and their respective untreated (UT) cells (H stands for HEK293 stable cells, L stands for LCA16 fibroblasts and R stands for iPSC-RPE). **(B)** Percentage of sequencing reads with substitutions as observed by deep sequencing analysis in BE-treated HEK^W53X^, Fibro^W53X^, iPSC-RPE^W53X/W53X,^ and their respective untreated cells. **(C)** Percentage of sequencing reads with total editing including substitutions and NHEJ as observed by deep sequencing analysis of optic cup gDNA from ABE8e treated Kcnj13^W53X/+^ mice (n=3 eyes) compared to a negative control Kcnj13^W53X/+^mouse treated only with PBS (n=1 eye). **(D)** Percentage of sequencing reads with only indels as observed by deep sequencing analysis in ABE8e treated Kcnj13^W53X/+^ (n=3 eyes) compared to a negative control Kcnj13^W53X/+^mouse treated only with PBS.

**Supplementary Figure 14:**
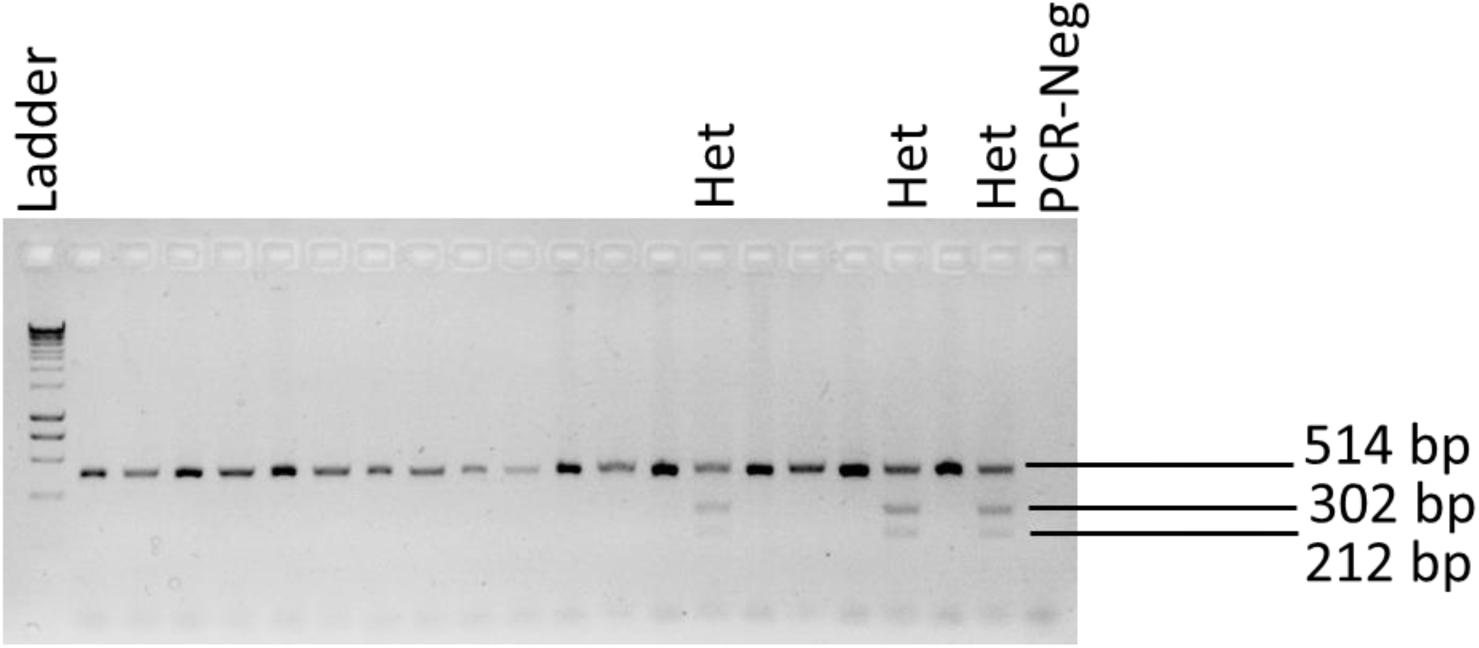
Agarose gel electrophoresis showing the differences in W53X heterozygous and WT mice. W53X mutation creates a restriction site for NheI; therefore, the W53X allele resulted in two (212 bp and 302 bp) fragments while the WT allele only one (514 bp) fragment.

